# Revisiting Reconstruction Likelihood: Variational Autoencoders for Biological and Biomedical Data Clustering

**DOI:** 10.64898/2026.04.09.717460

**Authors:** Andrej Korenić, Ufuk Özkaya, Abdulkerim Çapar

**Affiliations:** Institute for Physiology and Biochemistry “Jean Giaja”, Faculty of Biology, University of Belgrade, Serbia; Electrical and Electronics Engineering Department, Suleyman Demirel University, Isparta, Türkiye; Informatics Institute of Istanbul Technical University, Istanbul, Türkiye

**Keywords:** variational autoencoder, reconstruction likelihood, latent space clustering, out-of-distribution detection, biomedical data

## Abstract

**Background and Objective:** Variational Autoencoders (VAEs) offer a powerful framework for unsupervised anomaly detection and data clustering, often surpassing traditional methods. A core strength of VAEs lies in their ability to model data distributions probabilistically, enabling robust identification of anomalies and clusters through reconstruction likelihood — a stochastic metric providing a principled alternative to deterministic error scores.

**Methods:** We investigated how different VAE architectures, combining reconstruction likelihood with a learnable or data-driven prior, performed in a clustering task on a toy dataset such as MNIST. Results were verified using dimensionality reduction techniques like t-distributed Stochastic Neighbor Embedding (t-SNE) and Uniform Manifold Approximation and Projection (UMAP), alongside clustering algorithms such as k-means and Hierarchical Density-Based Spatial Clustering of Applications with Noise (HDBSCAN).

**Results:** The VAE’s encoder inherently maps data points into a latent space exhibiting discernible cluster structure, as evidenced by alignment with ground truth labels. While dimensionality reduction techniques (both t-SNE and UMAP) facilitated the application of clustering algorithms (k-means and HDBSCAN), these methods were primarily used to visualize and interpret the latent space organization.

**Conclusions:** This study demonstrates that VAEs effectively cluster data by implicitly encoding assignments in their latent representations. Determining cluster membership from encoder output, combined with reconstruction likelihood using semantic features, offers a principled approach for identifying typical samples and anomalies. Future research should focus on leveraging this inherent clustering capability of VAEs to enhance interpretability and facilitate clinical application.

## 1. Introduction

### 1.1. Clustering in biomedical research and challenges specific to biological data

Clustering is a fundamental unsupervised learning paradigm widely used across biological and biomedical research to discover inherent grouping structure in unlabeled data. [1–3] Examples include grouping cell states in singlecell omics, stratifying patients by multi-omics profiles, identifying phenotypic classes and changes in medical imaging, and detecting conformational substates in molecular simulations. [4, 5]

Despite its broad utility, clustering in biomedical contexts faces particular challenges: high dimensionality, strong noise and sparsity, mixed data types (e.g., continuous, count, categorical), technical and biological confounders, and frequent absence of clear, ground-truth cluster structure. [4] In a recent paper on clustering abnormal nuclear morphologies, the authors highlighted plainly the problem of clustering: “the output is simply a group of clusters without information as to what these clusters correspond to, thereby restricting its use in direct medical contexts.” [6] It is evident this complicates both the computational task of forming stable, reproducible clusters and the downstream clinical or biological interpretation required for translational use. [4] Moreover, despite advances in deep learning for medical image analysis, widespread clinical adoption is hindered by insufficient evidence of model reliability, which is critical in high-stakes medical decisions. [7]

### 1.2. Deep generative models as a representation-learning paradigm

Over the last decade, deep generative models — most prominently variational autoencoders (VAEs) and related autoencoder (AE) families — have emerged as a general representation-learning paradigm that naturally supports downstream clustering. VAEs learn nonlinear latent embeddings that compress high-dimensional features into lower-dimensional, often more interpretable manifolds. [8] These latent spaces can then be explored with standard clustering methods (e.g., Gaussian mixture models, k-means) or integrated with clustering objectives during training. In latter context, the network loss typically refers to the reconstruction loss of an AE or VAE and is crucial for initializing parameters and preventing cluster collapse. A separate loss, dubbed the clustering loss, encourages the model to learn representations that are conducive to grouping data into distinct clusters. [5]

A comprehensive review of combining neural networks with traditional clustering algorithms categorizes current deep clustering models into three groups based on network architecture and loss function characteristics: [5]

- AE-based deep clustering algorithms that use an AE as the core frame-work where network parameters are first initialized using reconstruction loss before introducing the clustering loss.
- Feed-forward networks trained solely by a specific clustering loss, referred to as direct cluster optimization; their performance depends on the robustness of the clustering loss.
- Models based on generative approaches such as VAEs, which can generate new samples while performing clustering; they inherit the disadvantages of AE-based algorithms and suffer from high computational complexity.

### 1.3. Latent space exploration, evaluation pitfalls and interpretability

In practice researchers frequently apply additional dimensionality reduction or visualization techniques to VAE latent spaces post-training in order to reveal and inspect putative clusters, and then must validate and interpret those clusters using biological annotation. [8] Regarding such approaches, it has been noted that “unsupervised deep learning on VAE models can autonomously learn biologically meaningful structures, many of which align with hand-crafted features previously defined” by researchers. [6]

A relatively recent review of deep-learning approaches to clustering in bioinformatics, conducted as these methods began to gain traction and be implemented in the field, documents consistent improvements in representational quality and clustering performance over classical pipelines (dimensionality reduction followed by distance-based clustering) especially in high-dimensional, noisy domains. [8] A key critique notes that distance-based clustering and standard benchmarks can mislead when datasets lack true cluster structure. Namely, partition measures may show strong agreement even for arbitrary groupings. This flaw undermines naïve benchmarking, urging careful reevaluation of clustering evaluation in biomedical data. [4]

Additionally, it was stressed out that incorporating established interpretability techniques would further deepen our understanding of the features driving these learned representations. [6] However, in this manuscript, we revisit a metric introduced now ten years ago — *reconstruction probability* (more exactly likelihood) — and note that its significance for biomedicine had already been highlighted, yet it remained overlooked and underappreciated. [7, 9–12]

### 1.4. Research objectives and paper organization

Our initial inquiry concerned whether clusters could be extracted directly from the latent space by using reconstruction likelihood. Specifically, after training a VAE, each new sample is mapped into the region of latent space that best matches it, effectively following learned clustering pathways. Moreover, by encoding two samples through the encoder, their representations can be compared; ideally, one would quantify the similarity or dissimilarity by estimating the probability that they belong to the same cluster without having to define clusters a priori. Finally, a VAE can construct an informative latent space so that, once labels are available, meaningful groups may emerge and be validated against those labels.

Subsequently, we examined which metrics are required to characterize the latent space and evaluate its reliability, essentially delineating the criteria that clustering methods must satisfy. Our second goal was therefore to provide an overview of these metrics from the perspective of comparing the performance of different VAE architectures.

The remainder of this manuscript is organized as follows: first, we establish the notation and explain how VAEs enable representation learning; next, we discuss reconstruction likelihood, its role and relevance in the VAEs under study; finally, we present a comparative performance evaluation of VAEs in clustering tasks on MNIST dataset and interpret the results with respect to biological and biomedical data.

## 2. Autoencoders and Variational Autoencoders

Autoencoders (AEs) and Variational Autoencoders (VAEs) are neural-network models that learn compact, informative representations of high-dimensional data — called **codings** or **latent variables**. An AE is a deterministic pair of multi-layer dense networks, an encoder and a decoder (architecturally MLPs), that maps an input to a low-dimensional latent vector and then reconstructs the input from that vector. Namely, with an AE we model each observation **x** ∈ 𝒳together with an associated latent variable **z** ∈ *Ƶ*. First, the encoder learns the representation of input **x** in a compressed format, mapping and transforming the data into an embedding **z**. Then the decoder tries to reconstruct 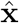 from **z** by minimizing the reconstruction loss between **x** and its corresponding reconstruction 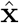. [8, 13, 14]

### 2.1. Generative models and inference networks

A VAE is a probabilistic graphical model that integrates variational inference with deep learning, providing a theoretically sound method for latent space learning (Fig. 1). The encoder outputs parameters of a distribution (typically Gaussian) over the latent space **z**, and the decoder samples from this distribution to generate reconstructions 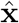. These representations are useful for dimensionality reduction, feature extraction, anomaly detection, and generative modeling. The underlying learning procedure can be interpreted as variational inference, closely related to the expectation-maximization algorithm. [15–17]

**Figure 1:**
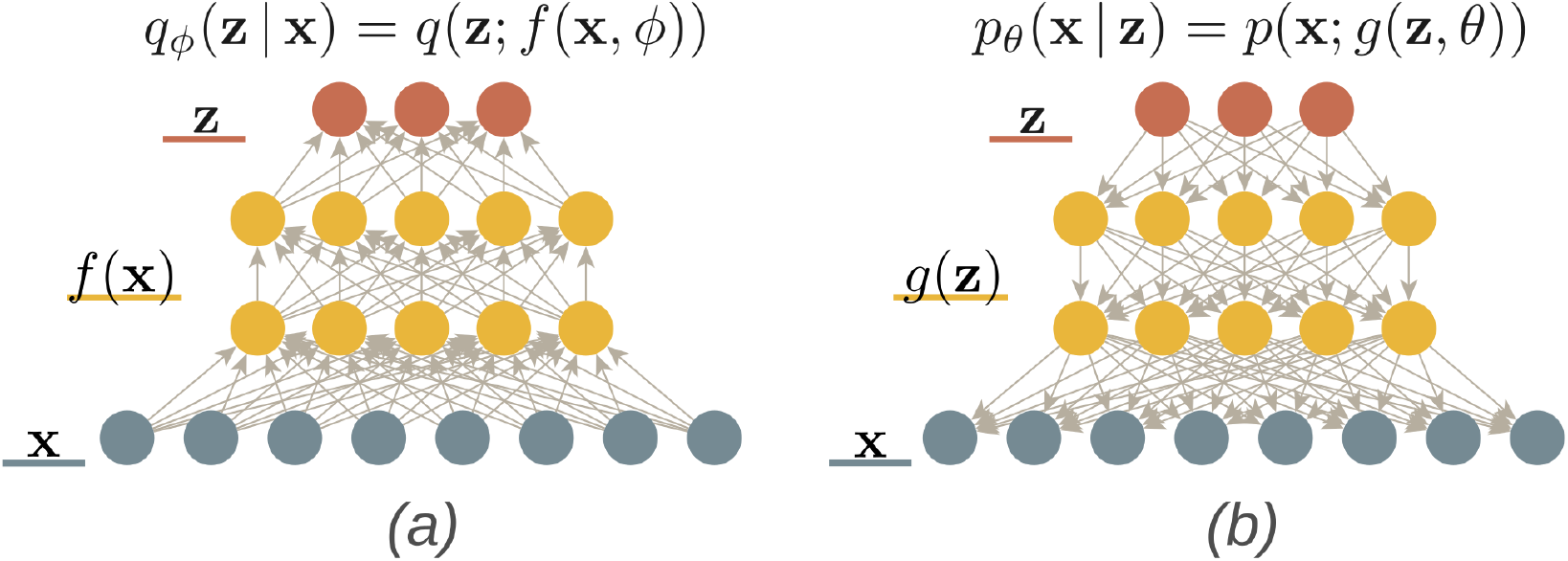
Adaptation of the schematic diagram of the encoder architecture (*left*) and decoder (*right*) as depicted in the work of An and Cho [9]. The colors of the neural network layers correspond to mathematical notation.

More specifically, the generative model (a top-down generative network), parameterized by *θ*, consists of a prior distribution *p*(**z**) and a likelihood *p*_*θ*_(**x** |**z**), which together define the joint distribution *p*_*θ*_(**x, z**) = *p*(**z**) *p*_*θ*_(**x**| **z**). The likelihood specifies the probability of observing **x** given a particular latent variable **z**, thereby determining how latent representations map to observable data. The prior *p*(**z**) is commonly a standard Gaussian 𝒩 (0, *I*) and it refers to the probability distribution over the latent variables **z** before observing any data **x**. It is a key component of the generative model, defining how latent variables are expected to be distributed in the absence of specific input data.

Practically, sampling **z** ∼*p*(**z**) and evaluating *p*_*θ*_(**x** |**z**) yields a reconstruction or, more specifically in the case of VAEs, a synthetic observation 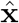. In other words, the decoder samples a latent code **z** from this distribution and maps it back to data space. This process specifies how data are generated from the latent space, revealing the generative properties of VAEs.

To obtain an approximation of a true posterior *p*(**z** |**x**), i.e., a distribution over latent variables **z** conditioned on a given observation **x**, we employ an inference network (a bottom-up recognition network; encoder) with parameters *ϕ*. Namely, this network defines an approximation *q*_*ϕ*_(**z** |**x**) to the true posterior *p*(**z** |**x**), since the latter is generally not analytically tractable. The encoder takes an input **x** and produces two outputs: a latent mean vector, denoted by *µ*_*ϕ*_(**x**) or *µ*_**z**_(**x**), and a log-variance vector, denoted by log 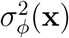 or log 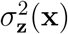. These parameters define the approximate posterior distribution *q*_*ϕ*_(**z** | **x**) = 𝒩 (*µ*_*ϕ*_(**x**), 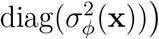. (For more details on standard VAE refer to Appendix A.1.)

Practically, for each pixel of an image, the decoder outputs a Bernoulli probability if the image is binary; for grayscale MNIST we replace that simple Bernoulli likelihood with a discretised logistic distribution (see Appendix A.3) so that the model can assign a meaningful probability mass to an integer value matching the 8-bit representation of each pixel intensity.

### 2.2. Training objective: evidence lower bound (ELBO)

Both the likelihood and the approximate posterior are parameterized by neural networks: the decoder network computes the parameters of *p*_*θ*_(**x** | **z**), while the encoder network produces the parameters of *q*_*ϕ*_(**z**| **x**). As we already said, the prior *p*(**z**) is fixed, typically a standard normal distribution, namely multivariate Gaussian denoted by 𝒩 (0, *I*), and does not depend on the data. In other words, VAEs utilize a probabilistic encoder to model the distribution of latent variables, allowing the system to account for the variability within the latent space through a sampling procedure. This contrasts with standard autoencoders, which model the latent variable itself. [9]

The training objective is a weighted sum of two terms. First, the Kullback– Leibler (KL) divergence between the approximate posterior *q*_*ϕ*_(**z** |**x**) and a unit Gaussian prior *p*(**z**) encourages the latent space to be smooth and well-structured: it penalises deviations from the standard normal, thereby preventing the encoder from collapsing into arbitrary, disconnected regions. Second, the reconstruction loss (computed as the negative log-likelihood of the observed values under the decoder’s distribution) forces the model to faithfully reproduce the input.

Together, these two components balance regularisation and data fidelity: the KL term shapes a latent manifold that is easy to sample from, while the reconstruction term ensures that samples drawn from this manifold can be decoded into realistic images that resemble the training set. In other words, during training, the VAE optimises for both terms simultaneously:

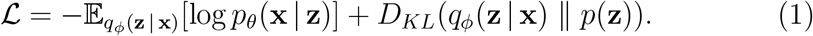

### 2.3. Reconstruction likelihood

#### 2.3.1. Origins and conflicting definitions

The work of An and Cho [9] on *reconstruction probability*, as they called it (but to be precise it is a **reconstruction likelihood**), introduced a principled method for anomaly detection using VAEs. They suggested computing the likelihood of data under the VAE model, thus treating low-probability observations as anomalies, providing a more robust alternative to simple reconstruction error by incorporating the inherent uncertainty (variance) of the VAE’s probabilistic framework.

However, their paper has been noted for inconsistent definition of *reconstruction probability* — initially defined as the expectation of the loglikelihood and later described as the expectation of the probability. This mismatch has led to divergent interpretations in subsequent research on whether one should “resample from the prior *p*(**z**) and evaluate *p*(**x** | **z**) to obtain a probability density of **x** […] or to derive useful outputs with the variational posterior *q*_*ϕ*_(**z** | **x**)” by averaging the log-likelihoods. [18, 19]

#### 2.3.2. Computing reconstruction likelihood

Namely, a straightforward (naïve) approach would be to sample multiple times from the latent space **z**, decode each sample to obtain a reconstruction 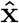, and obtain a multivariate normal distribution for every feature value (e.g., pixel intensity for images or signal amplitude for time-series). Other creative ideas included passing the decoded output through the encoder once again, hence relying on the error in encoding [20], or estimating the structure of a neighbourhood around the input point in latent space rather than reconstructing the point itself. [21] In these approaches, a coherent point in latent space is expected to generate a coherent decoder output with small variance, whereas a fuzzy or blurred point would produce a correspondingly fuzzy or blurred reconstruction with large variance.

As we already briefly mentioned above, in VAEs the decoder aims to reconstruct the original input data **x** from a given latent variable **z**. The likelihood of this reconstruction process is quantified by *p*_*θ*_(**x** |**z**), where *θ* represents the parameters of the decoder. This term, often referred to as the **reconstruction likelihood**, is typically maximized during training and is related to the overall objective function which also includes a regularization term based on the approximate posterior distribution *q*_*ϕ*_(**z** |**x**). Specifically, it is the Monte Carlo estimate of the expected log-likelihood of the data given the latent variables: 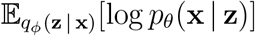. [9] (see also Supp. Fig. 1 and Appendix A.2 for more details.)

#### 2.3.3. Reconstruction likelihood across feature levels

All of the models we describe here and evaluate share a common feature: they effectively employ reconstruction likelihood. Although the work of An and Cho [9] specifically highlighted its importance for anomaly detection, it turns out that reconstruction likelihood will also be key for tasks such as recognizing data that does not belong to the initial training set, as well as for clustering on its own.

In fact, in the field of anomaly detection, an unsupervised framework has been already introduced that leverages representations automatically learned by a deep neural network to improve detection performance via subsequent clustering. Namely, the framework first trains an autoencoder and then applies a k-nearest neighbors (kNN) outlier detector, thereby simulating the identification of out-of-distribution (OOD) samples. [7, 22] Additionally, the authors of that study explored and detailed alternative methods for learning outliers in datasets as well as other anomaly detection techniques. [22]

Nevertheless, it has also been shown that employing such a criterion may be effective only for a specific dataset and not in more general scenarios. For instance, MNIST data can be surprisingly well reconstructed by a hierarchical VAE (HVAE) trained on Fashion-MNIST, confirming that the underlying cause of OOD failures is that highly generalizable low-level features can significantly contribute to estimated likelihoods, leading to poor performance in OOD detection [17]

More importantly, the key insight from the latter study is that in HVAEs one can evaluate reconstruction likelihood at different levels of the latent hierarchy by employing a likelihood-ratio score that compares reconstruction likelihoods across feature levels to detect OOD data. Therefore, advanced technique like likelihood-ratio score was proposed. This score aims to isolate high-level semantic features, allowing for better distinction between known (in-distribution) and unknown (out-of-distribution) data by reducing the influence of misleading low-level features. [17]

### 2.4. Importance Weighted Autoencoder (IWAE)

While a standard VAE draws one latent vector **z**_*i*_ per data point **x**_*i*_, an Importance-Weighted Autoencoder (IWAE) [23] generalises this by drawing *K* independent reparameterised samples from the approximate posterior (encoder) *q*_*ϕ*_(**z**_*i*_ |**x**_*i*_) for each observation **x**_*i*_.

Importance weighting is crucial because it lets the model exploit multiple latent draws per datum, reducing variance in the likelihood estimate and allowing a richer exploration of high-probability regions of the posterior. Consequently, IWAEs can often learn more expressive latent representations than standard VAEs. [23, 24] (For more details see Appendix B.)

### 2.5. VAEs with Gaussian mixture distributions

As we have shown so far, standard VAEs typically model the prior distribution of latent representations as a single multivariate Gaussian. However, a single Gaussian distribution excessively limits the expressive capacity of the model and is not conducive to obtaining cluster-friendly representations. Thus, attempts have been made to model the latent representation space with a Gaussian mixture distribution, and they show impressive performance, at least for the MNIST dataset that we explore here as a toy example. [5, 25]

#### 2.5.1. Variational mixture of posteriors (VampPrior)

The VampPrior [26] augments this setup with *K* learnable pseudo-inputs **u**_*k*_ of the same size 28×28 pixels. Real inputs contribute both reconstruction and KL terms to the ELBO, whereas pseudo-inputs influence only the KL term (through the *p*_*λ*_(**z**)) because they are never decoded (Supp. Fig. 1c).

Namely, in each training step every pseudo-input **u**_*k*_ is passed through the encoder just like a real datum, yielding posterior parameters (*µ*_*ϕ*_, *σ*_*ϕ*_). The prior over latent codes is defined as an average of these posteriors:

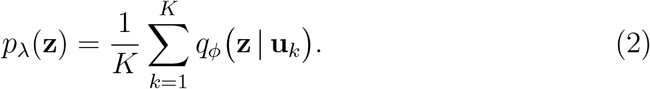

The gradients from the reconstruction loss on real data and the KL term jointly push the pseudo-inputs *q*_*ϕ*_(**z** | **u**_*k*_) toward “digit-like” prototypes. Each **u**_*k*_ therefore defines one Gaussian component in latent space; together they form a flexible, multimodal prior that replaces the standard isotropic Gaussian 𝒩 (0, *I*). Since each pseudo-input **u**_*k*_ is treated as an anchor that generates a posterior mean and variance, the collection of posteriors *q*_*ϕ*_(**z** |**u**_*k*_) forms a versatile mixture prior over **z**.

Sampling from this mixture tends to place latent codes in meaningful regions of the latent space, yielding sharper and more diverse reconstructions than sampling from a single unimodal Gaussian. After training, visualizing the learned pseudo-inputs usually reveals grayscale patterns that resemble typical MNIST digits — an empirical confirmation that they act as prototype anchors for the prior. (For more details see Appendix C.1.)

#### 2.5.2. Exemplar VAE

An Exemplar VAE also augments a standard VAE by replacing its fixed Gaussian prior with a data-driven prior built from latent encodings of real training samples (“exemplars”; see Supp. Fig. 1d) [27]. In a similar fashion to VampPrior, the Exemplar VAE uses two encoder passes:

1. *q*_*ϕ*_(**z** |**x**) denotes the variational posterior that encodes the target input **x**.
2. *r*_*ϕ*_(**z** | **x**_*n*_) denotes the variational posterior that encodes each exemplar **x**_*n*_ into a latent distribution that forms one component of the prior mixture.

During training we do not sample **z** from the encoder; instead we use the variational posterior *q*_*ϕ*_(**z** |**x**) (Supp. Fig. 1d). The prior in the ELBO becomes a mixture of the exemplar-induced posteriors, so that the KL divergence compares *q*_*ϕ*_(**z** | **x**) to this mixture prior defined by *r*_*ϕ*_(**z** | **x**_*n*_). The encoder *r*_*ϕ*_(**z** |**x**_*n*_) is trained jointly with *q*_*ϕ*_(**z** |**x**) by optimizing the Exemplar VAE’s ELBO, where both passes through the encoder share parameters.

In addition, Retrieval-Augmented Training (RAT) in the Exemplar VAE uses approximate nearest-neighbor search in latent space to identify the most relevant exemplars for a given input **x**. In other words, instead of using all available exemplars to form the prior, RAT selects only the nearest neighbors in latent space — those that are most influential for modeling **x**.

It should be noted that, in order to generate an image, we first select an exemplar **x**_*n*_ from the training set, encode it with the encoder *r*_*ϕ*_(**z** | **x**_*n*_) to obtain a latent distribution, sample a code **z** ∼ *r*_*ϕ*_(**z** | **x**_*n*_), and finally decode this code through *p*_*θ*_(**x** | **z**). The resulting observation is semantically related to the chosen exemplar **x**_*n*_. (For more details see Appendix C.2.)

## 3. Experiments: comparative evaluation of VAEs

Further in our study we performed a comparative analysis of the described VAEs (see Supp. Fig. 1) and measured their performance for clustering. In the following sections of this paper we provide a brief description of the methods used and present the results; for more information and details, see the Appendix and GitHub page (see footnotes section).

### 3.1. MNIST dataset

The MNIST dataset consists of grayscale images of handwritten digits, each represented as a 28 × 28 pixel image. We utilized the standard split of 60,000 training examples and 10,000 test examples. To facilitate model selection, we further partitioned the training set into a 50,000 examples training subset and a 10,000 examples validation subset. [23, 26, 27]

A key consideration in generative modeling with MNIST is the method of binarization. As noted by Burda et al. [23] the generative modeling literature exhibits inconsistencies regarding MNIST image binarization, with different choices potentially leading to considerably different log-likelihood values. We employed dynamic binarization during training, where each pixel value within an image is randomly converted to 0 or 1 using a Bernoulli sampling process.

### 3.2. Evaluation metrics

Table 1 succinctly maps our three core evaluation questions: recovery of true classes, internal coherence, and trade-off between compactness and separation, to the corresponding metrics. The table provides an at-a-glance roadmap for assessing clustering quality across both external and internal criteria. (For more details see Appendix E.)

**Table 1:** A quick summary of evaluation questions and the metrics that answer them.

| Question We Ask | Metrics That Help Us Answer It |
| --- | --- |
| Did we recover the true classes? | Accuracy (ACC), Adjusted Rand Score (ARI), V-measure Score (VMS), Fowlkes–Mallows Score (FMS), Adjusted Mutual Information Score (AMI) |
| Are the clusters internally coherent? | Silhouette Score (SS), Davies–Bouldin Index (DBI) |
| Is there a good trade-off between compactness and separation? | Calinski–Harabasz Index (CHI) |

#### 3.2.1. Reconstruction quality

VAEs are typically evaluated by tracking their evidence lower bound (ELBO) and related metrics. The ELBO (expressed as a negative loss) comprises both the reconstruction error (i.e., reconstruction likelihood) and the KL regularization term. In our study, we monitored the test marginal log-likelihood (LL) to assess how faithfully the VAEs compress data, thereby enabling direct comparison with prior work that employed similar evaluation protocols. [26, 27]

#### 3.2.2. Classification metrics

When ground-truth labels are available (as in classification problems) several extrinsic evaluation metrics can assess clustering quality by comparing the predicted cluster assignments to the known labels, effectively treating the clustering task as a classification problem. Table 2 summarizes the most widely used external metrics for comparing a clustering solution to known class labels. It links each metric to its core notion of agreement, thereby serving as a quick reference for selecting the appropriate diagnostic when evaluating how faithfully clusters recover true classes.

**Table 2:** These metrics quantify different aspects of agreement between a clustering solution and known class labels, capturing accuracy at both the sample and pairwise levels.

| Metric | What it Measures | How to Interpret the Result |
| --- | --- | --- |
| **Accuracy (ACC)** | Fraction of samples whose cluster assignment matches the true class label after an optimal one-to-one mapping between clusters and classes. | Values close to 1 indicate that clusters closely reproduce the known class labels; lower values suggest systematic mismatches. |
| **Adjusted Rand Index (ARI)** | Agreement between all pairs of samples based on whether they are assigned to the same or different clusters in both the clustering and the ground truth, corrected for chance. | Ranges from $-1$ to $1$; values near $1$ indicate strong agreement with ground truth, around $0$ correspond to random labeling, negative values mean worse-than-random. |
| **Adjusted Mutual Information (AMI)** | Amount of information shared between cluster assignments and true labels, normalized and adjusted for chance. | Ranges from $0$ to $1$; higher values indicate that cluster labels capture more information about the true classes and are robust to different numbers of clusters. |
| **Fowlkes–Mallows Index (FMI)** | Geometric mean of pairwise precision and recall, measuring how well pairs of points from the same class are clustered together and how well different classes are separated. | Ranges from $0$ to $1$; higher values indicate better preservation of pairwise relationships with few false merges or splits. |
| **V-measure Score (VMS)** | Harmonic mean of homogeneity (each cluster contains only one class) and completeness (all samples of a class are assigned to the same cluster). | Ranges from $0$ to $1$; higher values indicate clusters that are both pure and complete with respect to true classes. |

#### 3.2.3. Clustering metrics

Since the goal is to obtain separable latent clusters, internal cluster validity indices are natural choices (Table 3). [1–3, 6] If ground-truth labels are available, external metrics like adjusted rand score or classification accuracy are useful metrics to be reported. Accuracy in this context (matching ground-truth labels after optimal clustering) indicates that points can be reconstructed or assigned to the correct cluster. However, it does not reveal whether clusters are well separated from one another (low Davies–Bouldin index), whether each point is placed within a tight group (high Silhouette Score), or whether the overall cluster geometry is meaningful for downstream tasks such as anomaly detection, prototype selection, and visualization.

**Table 3:** These internal metrics evaluate cluster structure based solely on the data geometry, without reference to external ground-truth labels.

| Metric | What it Measures | How to Interpret the Result |
| --- | --- | --- |
| **Silhouette Score (SS)** | For each sample, compares the average distance to points in its own cluster ($a$) with the minimum average distance to points in the nearest other cluster ($b$). The score reflects how well samples fit within their assigned cluster relative to neighboring clusters. | Ranges from $-1$ to $+1$. Higher values indicate better-defined clusters; values near $+1$ imply strong separation, around $0$ suggest overlapping clusters, and negative values point to possible misassignment. |
| **Davies–Bouldin Index (DBI)** | Average similarity between each cluster and its most similar (i.e., closest) cluster, computed from within-cluster dispersion and between-cluster separation. | Lower values indicate better clustering: compact, well-separated clusters yield small DBI; larger scores suggest overlapping or poorly separated clusters. |
| **Calinski–Harabasz Index (CHI)** | Ratio of between-cluster dispersion to within-cluster dispersion, measuring how distinct clusters are relative to their internal variability. | Higher values indicate better-defined clusters. The index is most useful for comparing different clustering solutions on the same dataset rather than as an absolute quality measure. |

### 3.3. Dimensionality reduction (embedding)

t-Distributed Stochastic Neighbor Embedding (t-SNE) [28] and Uniform Manifold Approximation and Projection (UMAP) [29] are two widely used nonlinear dimensionality reduction techniques designed for visualizing high-dimensional data in 2D or 3D space. Both aim to preserve meaningful structures in the data, but they differ in their mathematical foundations and practical behavior. While t-SNE can reveal global structure, it does so indirectly: its primary objective is local similarity preservation. The global structure emerges as a byproduct, but distances between clusters may not be meaningful. (For more details see Appendix F.1.)

### 3.4. Clustering algorithms

We utilized two standard unsupervised clustering techniques: *(1)* k-means [30], which partitions data into a fixed number of spherical clusters, and *(2)* HDBSCAN [31], a density-based algorithm that discovers clusters of arbitrary shape while automatically filtering out noise. Both were used to assess the organization of VAE’s learned latent representations. (For more details see Appendix F.2.)

When clustering data, one typically expects the algorithm to produce cluster labels; however, a VAE does not do so by default, and additional dimensionality reduction techniques likewise fail to yield such output. Consequently, we relied on the ground-truth labels because MNIST dataset is intended for a classification task. In addition, both k-means and HDBSCAN, after completing clustering, assign arbitrary identifiers to the resulting clusters. Therefore, to evaluate accuracy we had to align these identifiers with the ground-truth labels. To achieve this alignment, we employed two strategies: *(1)* leave-one-out k-nearest neighbors (LOO-kNN) supervised classification to optimally map cluster labels to ground-truth labels, and *(2)* a heuristic approach that assigns each cluster the majority ground-truth class of its members. (For more details see Appendix F.3.)

## 4. Results

We compare five models — a standard VAE, IWAE with *K*=5 and *K*=50 importance samples, VampPrior (500 pseudo-inputs), and Exemplar VAE (approximate *k*-NN prior with *k*=10) — across three embedding spaces (raw 40-dimensional latent, t-SNE, and UMAP) and three clustering/classification methods (LOO-kNN, *k*-means, HDBSCAN). Boldface indicates the best value in each column across all five models.

### Log-likelihood estimation

Table 4 presents the test log-likelihood and ELBO for all five models, estimated via importance sampling with *S*=5000 samples. VampPrior achieves the best log-likelihood (−82.29), closely followed by Exemplar VAE (−82.31). IWAE (*K*=50) comes third (−82.88), demonstrating that the tighter importance-weighted bound translates into better density estimation compared to the standard VAE (− 84.45). All results are consistent with published values [26, 27].

**Table 4:** Test Log-Likelihood and ELBO for Five VAE Models on Dynamic MNIST. Log-likelihood is estimated via importance sampling with *S*=5000 samples. Boldface indicates the best (least negative) value.

| Model | Test NLL | Test ELBO |
| --- | --- | --- |
| VAE | −84.45 | 88.04 |
| IWAE ($K=5$) | −83.37 | 86.19 |
| IWAE ($K=50$) | −82.88 | 84.84 |
| VampPrior | **−82.29** | **85.30** |
| ExemplarVAE | −82.31 | 85.38 |

### Raw latent space

The VAE’s 40 latent dimensions already capture most of the generative factors: LOO-kNN classification accuracy reaches ≈0.96 for the standard VAE and ≈0.98 for VampPrior and Exemplar VAE (Table 5). Both structured priors consistently improve all clustering and classification metrics over the standard and IWAE models. Exemplar VAE achieves the best scores across all metrics in the raw latent space.

**Table 5:** LOO-kNN Classification Performance on VAE Latent Space (Raw)

| Model | ARI | AMI | SS | DBI | CHI | ACC | FMS | VMS |
| --- | --- | --- | --- | --- | --- | --- | --- | --- |
| VAE | 0.9133 | 0.8998 | 0.0306 | 3.3691 | 278.48 | 0.9602 | 0.9220 | 0.9000 |
| IWAE ($K=5$) | 0.8968 | 0.8838 | 0.0306 | 3.3708 | 267.87 | 0.9526 | 0.9072 | 0.8840 |
| IWAE ($K=50$) | 0.9080 | 0.8958 | 0.0310 | 3.3450 | 271.88 | 0.9579 | 0.9173 | 0.8960 |
| VampPrior | 0.9595 | 0.9491 | 0.1644 | 2.1572 | 886.67 | 0.9814 | 0.9636 | 0.9492 |
| ExemplarVAE | **0.9641** | **0.9533** | **0.3188** | **1.2743** | **2224.97** | **0.9835** | **0.9677** | **0.9533** |

*k*-means in the 40-dimensional latent space struggles to form tight clusters because Euclidean distance becomes less meaningful when many dimensions are noisy or redundant (Table 6). HDBSCAN fails to produce any clusters for models with a standard normal prior (VAE, IWAE), while VampPrior and Exemplar VAE yield meaningful clusters, with Exemplar VAE clustering 57.5% of the data at near-perfect accuracy (Table 7).

**Table 6:** K-Means Clustering Performance on VAE Latent Space (Raw)

| Model | ARI | AMI | SS | DBI | CHI | ACC | FMS | VMS |
| --- | --- | --- | --- | --- | --- | --- | --- | --- |
| VAE | 0.5450 | 0.6105 | 0.0314 | 2.9844 | 312.24 | 0.7237 | 0.6000 | 0.6262 |
| IWAE ($K=5$) | 0.5782 | 0.6207 | 0.0305 | 2.9726 | 299.85 | 0.7732 | 0.6207 | 0.6214 |
| IWAE ($K=50$) | 0.6223 | 0.6483 | 0.0345 | 2.9949 | 298.69 | 0.8048 | 0.6603 | 0.6489 |
| VampPrior | 0.5528 | 0.6711 | 0.1790 | 1.7349 | 1059.37 | 0.7076 | 0.6382 | 0.6835 |
| ExemplarVAE | **0.7514** | **0.8373** | **0.3030** | **1.3855** | **2093.83** | **0.8466** | **0.8069** | **0.8500** |

**Table 7:** HDBSCAN Clustering Performance on VAE Latent Space (Raw)

| Model | ARI | AMI | SS | DBI | CHI | ACC | FMS | VMS | % |
| --- | --- | --- | --- | --- | --- | --- | --- | --- | --- |
| VAE | — | — | — | — | — | — | — | — | — |
| IWAE ($K=5$) | — | — | — | — | — | — | — | — | — |
| IWAE ($K=50$) | — | — | — | — | — | — | — | — | — |
| VampPrior | 0.7524 | 0.8873 | 0.2981 | 1.1951 | 865.35 | 0.8005 | 0.8180 | 0.8877 | 39.7 |
| ExemplarVAE | **0.9975** | **0.9958** | **0.4684** | **0.8726** | **2326.52** | **0.9986** | **0.9978** | **0.9958** | **57.5** |

### t-SNE embedding

t-SNE compressed the latent space into two dimensions while preserving local neighborhood structure, making cluster boundaries clearer for both density-based (HDBSCAN) and centroid-based (*k*-means) methods. LOO-kNN accuracy remains high (≈0.96–0.98; Table 8), while SS and DBI improve dramatically. *k*-means benefits substantially from dimensionality reduction, with Exemplar VAE achieving the best metrics across the board (Table 9). HDBSCAN now clusters all five models successfully, with VampPrior and Exemplar VAE reaching ≈0.98 accuracy at 97–99% coverage (Table 10).

**Table 8:**
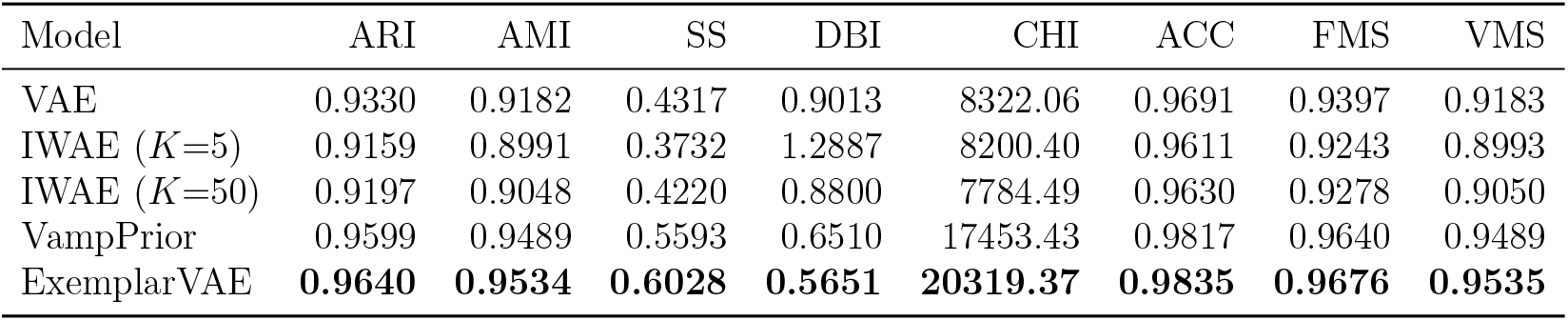
LOO-kNN Classification Performance on t-SNE Embedding of VAE Latent Space.

**Table 9:** K-Means Clustering Performance on t-SNE Embedding of VAE Latent Space.

| Model | ARI | AMI | SS | DBI | CHI | ACC | FMS | VMS |
| --- | --- | --- | --- | --- | --- | --- | --- | --- |
| VAE | 0.8425 | 0.8461 | 0.4832 | 0.6813 | 14254.53 | 0.9239 | 0.8584 | 0.8463 |
| IWAE ($K=5$) | 0.7121 | 0.7659 | 0.4479 | 0.7456 | 13092.60 | 0.8315 | 0.7412 | 0.7663 |
| IWAE ($K=50$) | 0.8372 | 0.8378 | 0.4603 | 0.7296 | 13140.37 | 0.9220 | 0.8536 | 0.8381 |
| VampPrior | 0.8221 | 0.8872 | 0.5399 | 0.6512 | 17884.96 | 0.8900 | 0.8760 | 0.9029 |
| ExemplarVAE | **0.9303** | **0.9297** | **0.6167** | **0.5337** | **23004.08** | **0.9675** | **0.9373** | **0.9298** |

**Table 10:** HDBSCAN Clustering Performance on t-SNE Embedding of VAE Latent Space.

| Model | ARI | AMI | SS | DBI | CHI | ACC | FMS | VMS | % |
| --- | --- | --- | --- | --- | --- | --- | --- | --- | --- |
| VAE | 0.5015 | 0.8008 | 0.3191 | 0.7732 | 5134.21 | 0.6816 | 0.6273 | 0.8011 | 89.8 |
| IWAE ($K=5$) | 0.8570 | 0.9147 | 0.5204 | 0.6091 | 13244.50 | 0.9109 | 0.8724 | 0.9149 | 78.6 |
| IWAE ($K=50$) | **0.9547** | **0.9459** | 0.5230 | 0.6639 | 13719.81 | 0.9790 | 0.9593 | 0.9461 | 82.5 |
| VampPrior | 0.9546 | 0.9456 | 0.5745 | 0.6229 | 21041.16 | 0.9791 | 0.9592 | 0.9457 | **99.0** |
| ExemplarVAE | 0.9313 | 0.9359 | **0.5833** | **0.5430** | **21415.42** | **0.9839** | **0.9685** | **0.9557** | 97.9 |

### UMAP embedding

UMAP produced an embedding space with even clearer geometric structure: SS, DBI, and CHI all improve compared to t-SNE, while classification metrics remain comparable (Table 11, Table 12, Table 13). HDBSCAN on UMAP achieved the highest overall coverage (97– 99.8%) with strong accuracy. VampPrior produced the best HDBSCAN results on UMAP, while Exemplar VAE dominated *k*-means metrics. All t-SNE and UMAP runs produced CHI values in the thousands to hundreds of thousands, far surpassing the raw 40-dimensional latent space, confirming that the low-dimensional embeddings create a high-contrast clustering structure (Supp. Fig. 2).

**Table 11:** LOO-kNN Classification Performance on UMAP Embedding of VAE Latent Space.

| Model | ARI | AMI | SS | DBI | CHI | ACC | FMS | VMS |
| --- | --- | --- | --- | --- | --- | --- | --- | --- |
| VAE | 0.9287 | 0.9146 | 0.6311 | 0.5085 | 25891.20 | 0.9672 | 0.9359 | 0.9147 |
| IWAE ($K=5$) | 0.9226 | 0.9076 | 0.6157 | 0.5423 | 25073.62 | 0.9643 | 0.9304 | 0.9078 |
| IWAE ($K=50$) | 0.9254 | 0.9119 | 0.6574 | 0.4593 | 28619.58 | 0.9657 | 0.9329 | 0.9120 |
| VampPrior | 0.9494 | 0.9393 | 0.7758 | **0.3328** | **86161.34** | 0.9768 | 0.9544 | 0.9394 |
| ExemplarVAE | **0.9578** | **0.9474** | **0.7794** | 0.3723 | 41742.84 | **0.9806** | **0.9620** | **0.9475** |

**Table 12:** K-Means Clustering Performance on UMAP Embedding of VAE Latent Space.

| Model | ARI | AMI | SS | DBI | CHI | ACC | FMS | VMS |
| --- | --- | --- | --- | --- | --- | --- | --- | --- |
| VAE | 0.7849 | 0.8510 | 0.6087 | 0.5824 | 29907.31 | 0.8674 | 0.8416 | 0.8660 |
| IWAE ($K=5$) | 0.7564 | 0.8283 | 0.5818 | 0.5776 | 19536.31 | 0.8394 | 0.7997 | 0.8359 |
| IWAE ($K=50$) | 0.9121 | 0.9015 | 0.6705 | 0.4465 | 36346.23 | 0.9593 | 0.9209 | 0.9016 |
| VampPrior | **0.9431** | **0.9347** | 0.7883 | 0.3141 | 148902.79 | **0.9738** | **0.9488** | **0.9349** |
| ExemplarVAE | 0.9262 | 0.9260 | **0.8256** | **0.2543** | **214159.27** | 0.9656 | 0.9337 | 0.9262 |

**Table 13:** HDBSCAN Clustering Performance on UMAP Embedding of VAE Latent Space.

| Model | ARI | AMI | SS | DBI | CHI | ACC | FMS | VMS | % |
| --- | --- | --- | --- | --- | --- | --- | --- | --- | --- |
| VAE | 0.9271 | 0.9172 | 0.6590 | 0.4592 | 43569.65 | 0.9701 | 0.9414 | 0.9228 | 97.1 |
| IWAE ($K=5$) | 0.7856 | 0.8666 | 0.6197 | 0.4862 | 21231.50 | 0.8561 | 0.8160 | 0.8668 | 97.7 |
| IWAE ($K=50$) | 0.9262 | 0.9137 | 0.6781 | 0.4382 | 37327.03 | 0.9659 | 0.9336 | 0.9138 | 98.7 |
| VampPrior | **0.9475** | **0.9383** | **0.7892** | **0.3122** | **149725.41** | **0.9759** | **0.9528** | **0.9384** | **99.8** |
| ExemplarVAE | 0.8224 | 0.8920 | 0.7316 | 0.3709 | 88790.02 | 0.8910 | 0.8826 | 0.9164 | 99.2 |

**Figure 2:**
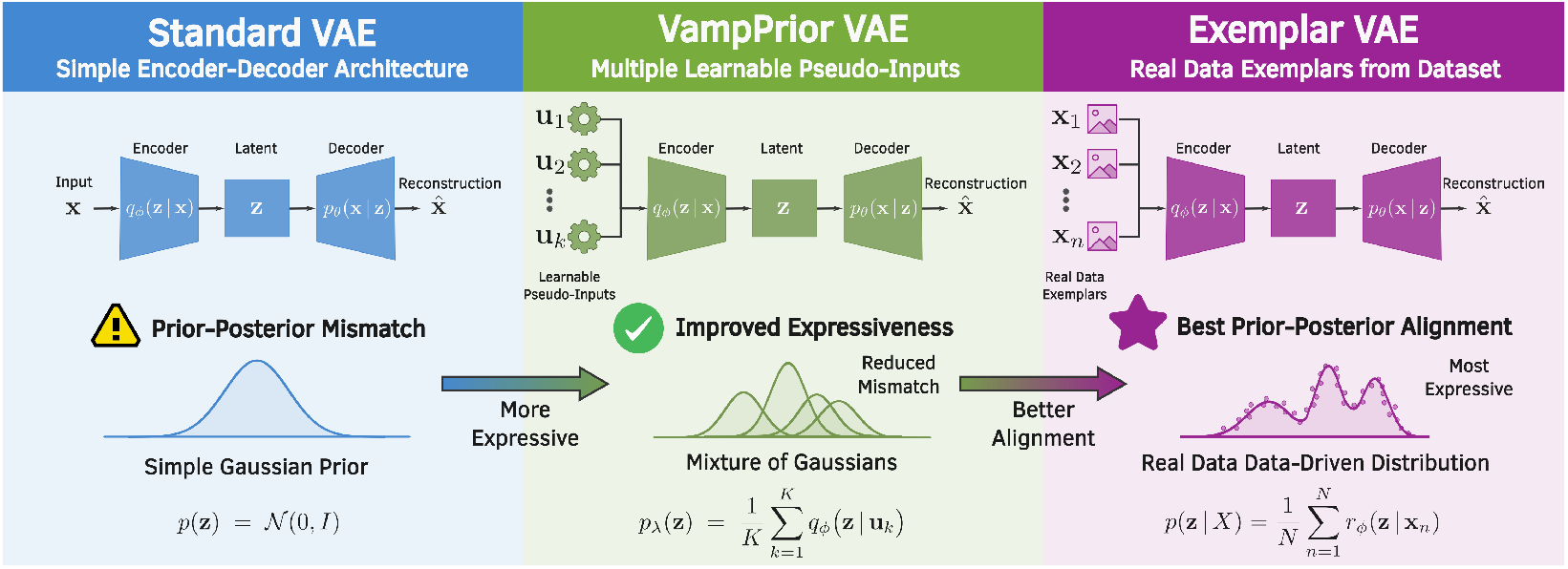
A schematic comparison of the performance of a standard VAE (*left*), VampPrior VAE (*middle*) and Exemplar VAE (*right*). Our visual representation follows “the line of thinking that the prior is a critical element to improve deep generative models, and in particular VAEs,” as emphasized by Tomczak and Welling [26].

## 5. Discussion

In our view, An and Cho [9] emphasized two pivotal and intertwined ideas in their paper that subsequent algorithms have leveraged: *(1)* a probabilistic decoder that outputs both mean and variance parameters rather than assuming a fixed variance; and *(2)* multiple resampling from the latent space to obtain tighter and more precise estimates of the likelihood.

By using these decoder-derived parameters, the likelihood of the original data given those parameters can be calculated, denoted **reconstruction likelihood**. In our work we aimed to demonstrate that a VAE, when adapted to use reconstruction likelihood and a variational mixture of posteriors, already supports meaningful clustering (hence, there is no need to rely on an external clustering method, post-festum in a sense). In other words, the VAE itself performs clustering because labels are not used during training. When we wish to assess how MNIST digits are grouped using their true labels, we can employ the same evaluation metrics used for classification and confirm the results.

Based on reconstruction likelihood, An and Cho concluded that the VAE learned the simple structure of digit 1 (a single vertical stroke) from other parts of the dataset. For instance, as they explained, a vertical stroke appears in nearly every digit: centrally in 4, on the right side in 7 and 9 when written rigidly without much curvature. [9, 32]

What this body of work presented so far demonstrates in paradigm terms is threefold. First, the VAE clustering paradigm is modality-agnostic: the same conceptual pipeline — learn a compressed latent representation with a generative model, then identify group structure in that latent space — has been successfully applied to diverse data types (single-cell and bulk transcriptomics, chromatin accessibility, metabolomics, protein structural ensembles, and imaging). Second, success depends critically on the design of the latent representation (architectural choices, reconstruction and regularization losses, probabilistic assumptions), on how confounders are controlled or removed, and on the choice of clustering mechanism used on the learned embedding. Third, although VAEs and related models often recover latent axes that align with biologically meaningful signals, these embeddings remain abstract and require careful downstream annotation, statistical validation, and confounder checks before any translational claims can be made. [33]

### 5.1. Is decoder even necessary?

A recent study reported that, for clustering tasks, the decoder used to reconstruct the original input becomes largely redundant once training concludes. Consequently, the authors demonstrated, both mathematically and empirically, that minimizing reconstruction error (the conventional objective of decoders) is equivalent to maximizing a lower bound on the mutual information (MI) between the input and its learned representation. [5] However, another study that also employed MI highlighted that a decoder is necessary because the lack of the decoder in the training process may cause the embedded feature space to collapse; i.e., using a decoder during training preserves the local structure from the original data in its embedded representations. [34]

We have already emphasized that there are also brilliant approaches which, instead of modeling reconstruction of the input, model the neigh-borhood of input information; these methods likewise yield strong results on the MNIST dataset. [21, 35, 36] The context-encoding VAE (ceVAE) combines context encoders with VAEs to address shortcomings in traditional VAE anomaly detection. Namely, ceVAE use both a better-calibrated reconstruction error and KL-divergence deviations, so it can be steered toward learning more discriminative, semantically rich features. [37] Likewise, other advanced VAE architectures, such as Nouveau VAE (NVAE) [38] and Ladder VAE (LVAE) [39], move beyond a single latent variable to incorporate partitioned groups of variables arranged hierarchically. Such hierarchical structures “enable the model to capture global long-range correlations at the top of the hierarchy and local fine-grained dependencies in lower groups”. [38] (For more comprehensive review Tschannen et al. [40].)

To move beyond the limitations of a standard multivariate Normal distribution, several approaches (SVAE [41], JointVAE [42], VQ-VAE [43], Vamp-Prior [26], and Exemplar VAE [27]) propose more structured priors in VAEs to improve interpretability, facilitate clustering, and enhance generative performance. These ‘meta-priors’ are intended to reflect general assumptions about data, such as their hierarchical organization or clusterability. [40]

Given our verification and experimental support for the clustering performance of VampPrior and Exemplar VAE, we are confident in their demon-strated results (see Fig. 2 and Supp. Fig. 2). Achieving comparable or slightly lower results with other methods confirms that these two models can learn superb latent representations and accurately determine reconstruction likelihood. Furthermore, reconstructing data and generating new samples should be desirable features; the model should allow exemplars to be reconstructed since they can serve as anchor points in the latent space while also determining data variability around each input feature.

### 5.2. On obtaining probability values

Clustering algorithms, particularly in biomedical applications, face several significant challenges, primarily stemming from the inherent complexity of biological data and the limitations of current evaluation methods. Many biological datasets may not possess intrinsic distance-based cluster structures, yet algorithms are often designed to ‘force’ a division, potentially leading to arbitrary and misleading labels. [4]

In the absence of clear labels, researchers rely on unsupervised quality measures like the Silhouette Score or Davies–Bouldin index. However, these measures are inherently biased, as they primarily evaluate how well an algorithm fits its own assumptions (e.g., spherical clusters) rather than reflecting actual biological reality. [4] Therefore, uncertainty quantification plays a crucial role in enhancing clinician trust in automated clusters and machine learning models, particularly in medical applications. [1, 3, 7]

The motivation for using reconstruction likelihood is largely sound — when applied correctly it offers a superior alternative to raw error for heterogeneous data. Nonetheless, the assertion that threshold selection is objective appears tenuous. [9] Namely, by employing log-likelihoods, we indeed obtain a more principled anomaly score that is invariant to the scale of different variables and yields thresholds that transfer better. In other words, reconstruction likelihoods address this threshold issue, making thresholds more objective and understandable.

Moreover, in anomaly detection, what we are typically interested in is indeed a reconstruction likelihood — that is, how “typical” a sample is under the learned distribution. However, this approach works as long as we recognize that the threshold parameter (analogous to a statistical *α* for testing *p*-value) must be tuned empirically, for example on validation data, to control the false-positive rate. For genuine statistical interpretability, one should rely on likelihood ratios, [17] cumulative distribution functions, or quantile-based thresholds. [11]

In addition, we must consider that VAEs trained on a single dataset can reconstruct data from other datasets; as we have already mentioned, for example, an HVAE trained on FashionMNIST can reconstruct digits from MNIST. Therefore, it is advisable to use the reconstruction likelihood evaluated with higher-level semantic features when clustering biological and biomedical data. [17]

## 6. Conclusion

In this study we showed that VAEs employing a reconstruction likelihood and a variational mixture posterior as a prior could be successfully used for clustering biological and biomedical data, as evidenced by their performance on MNIST toy example. The latent space warrants further investigation with additional dimensionality reduction methods such as t-SNE and UMAP, and clustering techniques such as k-means and HDBSCAN. However, we would like to emphasize that the VAE’s primary contribution lies elsewhere. Of exceptional importance is that, by passing the data through the encoder, one can obtain the location of these data points in the latent space and group them around prominent exemplars, while a reconstruction likelihood evaluated with higher-level semantic features ensures that the VAE recognizes whether it is observing data similar to those used for training or an OOD data.

## CRediT authorship contribution statement

**Andrej Korenić**: Conceptualization, Methodology, Software, Validation, Formal analysis, Resources, Writing - Original Draft, Writing - Review & Editing, Visualization. **Ufuk Özkaya**: Conceptualization, Methodology, Resources, Writing - Review & Editing, Supervision. **Abdulkerim Çapar**: Conceptualization, Methodology, Resources, Writing - Review & Editing, Supervision.

## Acknowledgements

The authors would like to thank Michele De Vita for his assistance in understanding the work of An and Cho, as well as his efforts to implement *reconstruction probability*. His GitHub code and insightful discussions were instrumental in structuring this study.

## Declaration of AI-assisted technologies in the manuscript preparation process

To enhance the language and readability of this manuscript, the authors used GPT OSS 20B employed in a local environment to ensure data privacy. The authors carefully reviewed edits suggested by the AI model, retaining full responsibility for the accuracy and integrity of the final manuscript.

## Supplementary Figures

**Supplementary Figure 1:**
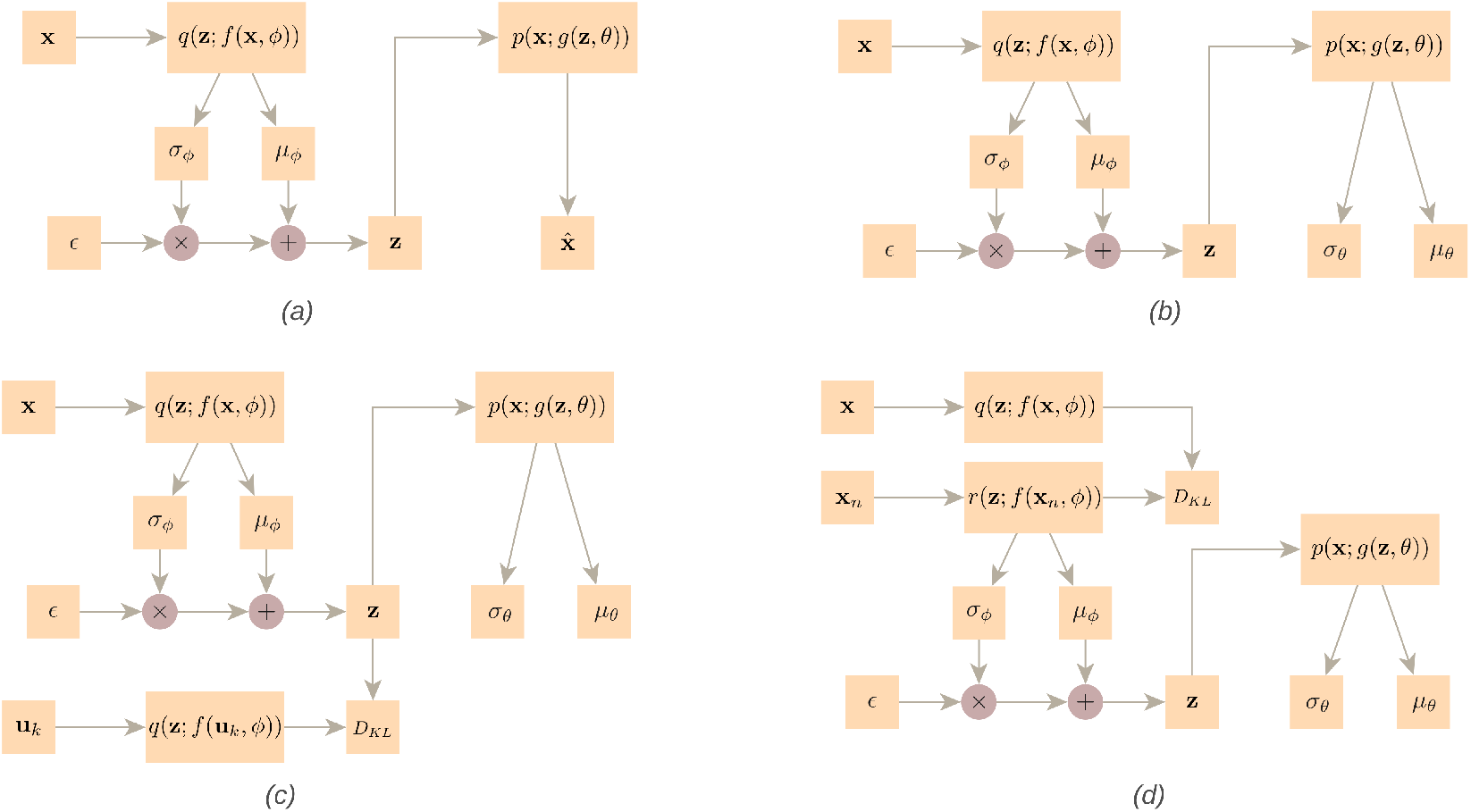
Diagrammatic representations of the various VAE architectures evaluated in this study: *(a)* a standard VAE with a decoder outputting only the reconstructed input value 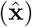, *(b)* a VAE whose decoder outputs both *µ*_*θ*_ and *σ*_*θ*_ as described in the work of An and Cho [9], *(c)* VampPrior [26], and *(d)* Exemplar VAE [27]. Encoder and decoder notation follows the convention in Figure 1. The reparameterization trick shown in equations A.6 and B.2 is used for sampling from the latent space; *ϵ* is a random noise variable drawn typically from a standard normal distribution. *D*_KL_ node denotes what goes into calculating Kullback–Leibler divergence. The pseudo-inputs **u**_*k*_ serve primarily to structure the prior, whereas the exemplars **x**_*n*_ are used both to structure the prior and to sample the latent space before the decoder reconstructs the input **x**.

**Supplementary Figure 2:**
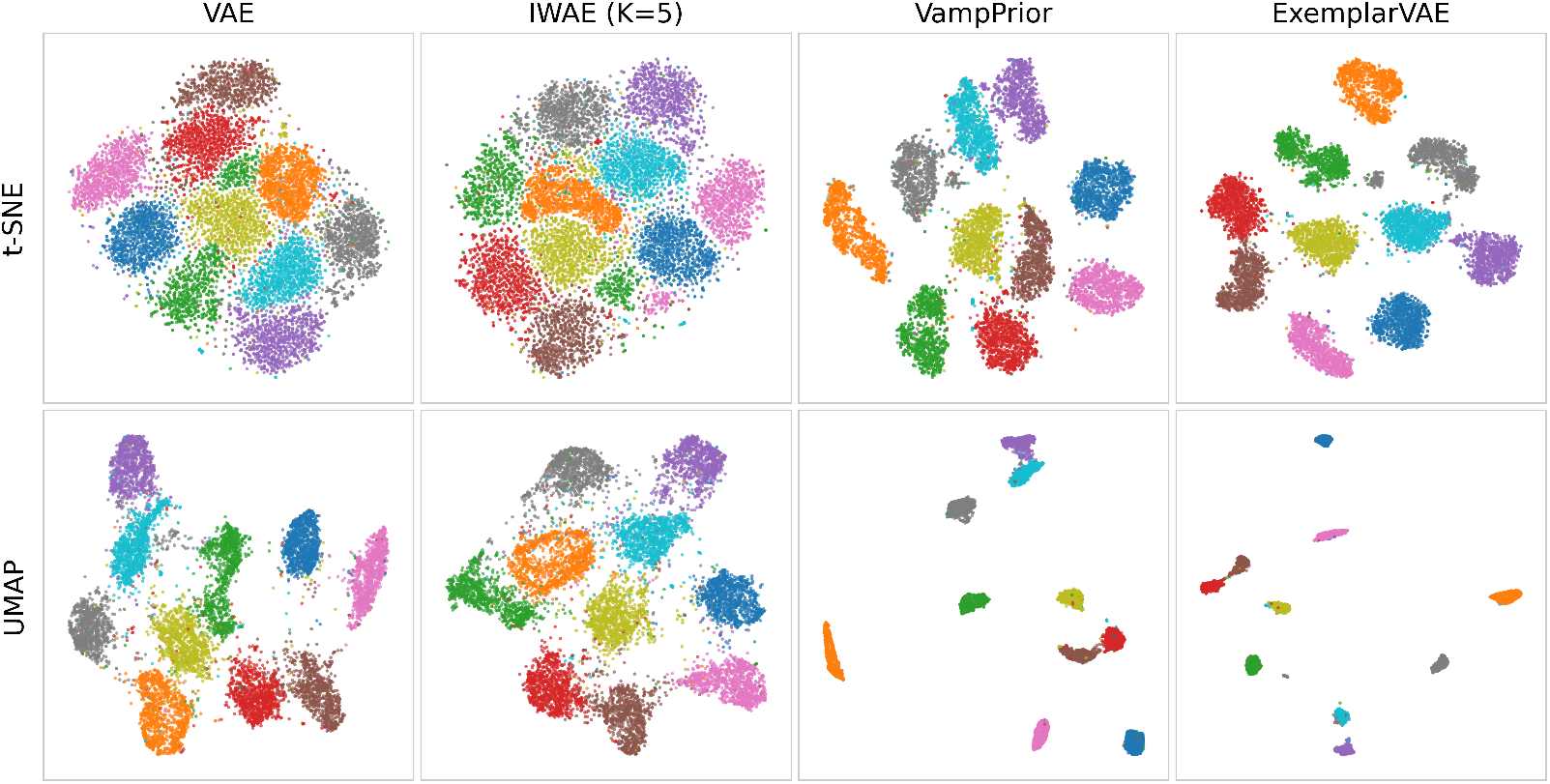
Visualization of the latent space using t-SNE (*top row*) and UMAP (*bottom row*), following dimensionality reduction. These visualizations suggest that VampPrior produces better-separated clusters than the standard VAE in both cases. Exemplar VAE visually provides just a slight improvement over VampPrior, although metrics provide stronger support for this conclusion. For comparison see Roellinger et al. [6], Norouzi et al. [27].

## Appendix

### A. Standard variational autoencoder

#### A.1. VAE formal definitions

A Variational Autoencoder (VAE) models the conditional distribution of the observed data **x** given a latent variable **z** as *p*_*θ*_(**x**| **z**). Using mean squared error (MSE) as the reconstruction loss in VAEs is theoretically justified when we assume that decoder models the likelihood *p*_*θ*_(**x** |**z**) as a multivariate Gaussian distribution with fixed (diagonal) covariance:

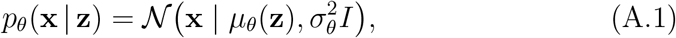

where *µ*_*θ*_(**z**) is the decoder output (the reconstructed mean), 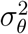 is a fixed scalar variance independent of **z** (i.e., hyperparameter, not learned), and *I* is the identity matrix (independent pixel noise). This assumption is standard for real-valued data.

The reconstruction term in the VAE objective is the expected negative log-likelihood:

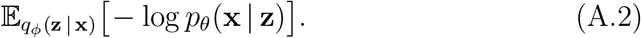

For the Gaussian likelihood above, the negative log-likelihood is:

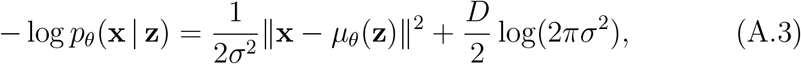

where *D* is the dimensionality of data **x**.

Since *σ*^2^ is fixed, the second term is constant with respect to the model parameters (i.e., w.r.t. *θ*), so the only trainable component in the likelihood term is the squared error ∥**x** −*µ*_*θ*_(**z**) ^2^∥. Therefore, minimizing the negative log-likelihood is equivalent (up to a constant scaling factor) to minimizing the sum of squared errors, while dividing by *D* yields the MSE.

In practice, the decoder’s output variance is often treated as a hyperparameter rather than being learned by the decoder. If variance were learned, maximum likelihood estimation could drive it toward zero to make the likelihood arbitrarily large, causing overfitting and poor latent representations. Thus, fixing it prevents this collapse and ensures a balance between reconstruction and the KL regularization term.

VAEs are built on variational inference, where the goal is to maximize the Evidence Lower Bound (ELBO) for each observed data point **x**. The ELBO serves as a tractable lower bound on the log-likelihood log *p*_*θ*_(**x** |**z**), enabling efficient optimization in the presence of latent variables **z**. The objective is defined as:

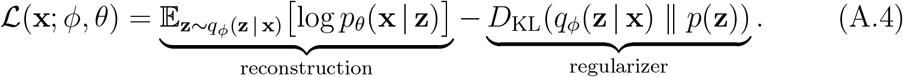

During training, the negative ELBO is minimized as the effective loss function. The first term, 𝔼_*q*(**z** | **x**)_[log *p*_*θ*_(**x z**)], is the expected reconstruction likelihood, which measures how likely the input **x** is under the decoded distribution *p*_*θ*_(**x** | **z**). Under the fixed-variance Gaussian assumption:

- Maximizing the reconstruction term is equivalent to minimizing MSE, and this term encourages the model to generate outputs that closely resemble the input data.
- The Kullback–Leibler (KL) divergence term regularizes the latent space independently of the reconstruction metric, which enforces similarity between the approximate posterior distribution *q*_*ϕ*_(**z** |**x**) — produced by the encoder — and the predefined prior *p*(**z**), typically set to a standard multivariate Gaussian 𝒩 (0, *I*).

Because the expectation is generally intractable, in practice we approximate it by drawing one or more samples (**z**^(*s*)^ ∼*q*_*ϕ*_(**z** | **x**)) and averaging:

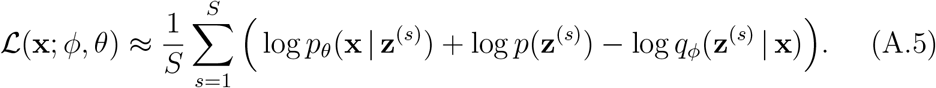

In standard VAE implementations the reconstruction term is estimated by sampling (using *S* = 1 for efficiency) while the KL term is computed analytically (e.g. when both are Gaussians). That hybrid reduces variance and computation. The expectation 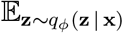 is approximated using a single sample via the reparameterization trick (Monte Carlo approximation):

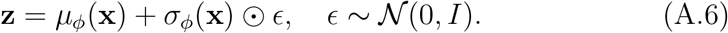

The encoder network defines a probabilistic mapping from the input space to the latent space by specifying the parameters of an approximate posterior distribution. In this work, we adopt a multivariate Gaussian with a diagonal covariance structure for the encoder output:

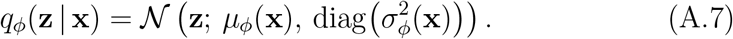

Here, *µ*_*ϕ*_(**x**) denotes the mean vector and 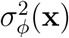 represents the vector of standard deviations, both parameterized by a neural network with weights *ϕ*. The resulting covariance matrix is constrained to be diagonal, implying conditional independence among the latent dimensions given the input. This assumption enables computational efficiency and facilitates the derivation of closed-form expressions for the KL divergence.

Under the assumption that the prior *p*(**z**) is a standard normal distribution 𝒩 (0, *I*) and the approximate posterior *q*_*ϕ*_(**z** |**x**) is a diagonal Gaussian, the KL divergence between these two distributions admits an analytical solution. This avoids the need for stochastic estimation and improves training stability.

The expression is given by:

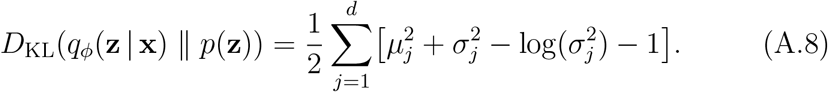

where *µ*_*j*_ = [*µ*_*ϕ*_(**x**)]_*j*_ and *σ*_*j*_ = [*σ*_*ϕ*_(**x**)]_*j*_ denote the *j*-th components of the mean and standard deviation vectors, respectively; and 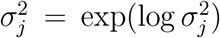. This widely used formula arises from properties of multivariate Gaussians with diagonal covariances and ensures that gradients can be efficiently back-propagated through the latent variable sampling process when combined with the reparameterization trick.

The KL term depends only on encoder parameters 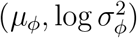 and the prior. It does not involve the observation variance 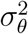, whether fixed or learnable. Fixing 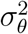 simplifies training and reduces the number of parameters, so that the decoder only needs to predict the mean *µ*_*θ*_(**z**) and the variance 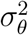 is treated as a hyperparameter, often set to 1 or tuned empirically.

Moreover, the KL divergence term plays a critical role in structuring the learned latent space. When the encoder produces posterior distributions *q*_*ϕ*_(**z** | **x**) that significantly deviate from the unit Gaussian prior — such as exhibiting large mean values (| *µ*_*j*_|≫ 0) or high variance (*σ*_*j*_≫ 1) — the KL penalty increases, discouraging such configurations. Consequently, the optimization process balances two competing goals:

- Faithful data reconstruction, driven by maximizing the expected log-likelihood under the decoder,
- Latent space regularization, achieved by minimizing the divergence from the prior.

This trade-off encourages the encoder to learn compact, smooth, and well-organized representations that are both informative for reconstruction and amenable to sampling, thereby supporting effective generative modeling. As a result, the latent space approximates a continuous, structured manifold where semantically similar inputs are embedded in proximity.

#### A.2. Reconstruction likelihood: decoder outputs variance

Firstly, we can replace the single decoder output, denoted by *µ*_*θ*_(**z**) or 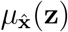, [9, 19] with two outputs:

- *µ*_*θ*_(**z**) — predicted mean (same shape as **x**),
- logvar_*θ*_ (**z**) — predicted log-variance (same shape as **x** for heteroscedastic noise).

We use log-variance as the canonical parameterization because it guarantees positivity of *σ*^2^ after exponentiation and is numerically stable. Under *p*_*θ*_(**x** | **z**) = 𝒩(**x** | *µ*_*θ*_(**z**), 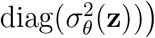 with 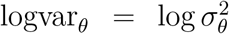, the per-datapoint negative log-likelihood becomes (up to standard constants):

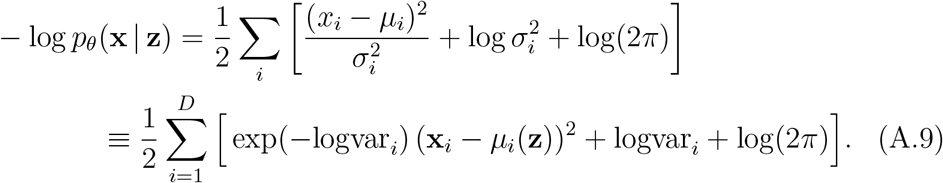

The reconstruction loss now includes a weighted MSE term, namely 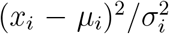, and a log-variance penalty, 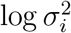. Learnable variance can improve flexibility but often complicates training (e.g., variance collapse / shrinkage; see [10]). If used, regularization on log 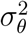 is usually required (see HardTanh, A.4). Because log(2*π*) is a constant, it can be omitted when comparing losses.

##### A.2.1. Variance, logits and uncertainty

In an AE, the minimization of the reconstruction loss (typically mean squared error, MSE, or binary cross-entropy, BCE) is employed to improve reconstruction quality. However, when the same principle is applied to a standard VAE, the decoder learns the mean values while keeping the variance fixed. More specifically, the Kullback–Leibler (KL) divergence term enforces that the learned latent distribution approximates a prior (usually a standard normal distribution) thereby promoting smoothness and interpretability in the latent space. However, because the variance is fixed, the VAE has no incentive to learn a pixel-wise uncertainty estimate. The only thing it can adjust is the mean, denoted by *µ*_*θ*_(**z**) or 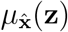 (see [9, 19] and Appendix A.2). In other words, minimizing the MSE is equivalent to maximizing the log-likelihood of the data when the errors are assumed to follow a Gaussian distribution, particularly when *σ*^2^ = 1.

On the other hand, in a VAE decoder, logits refer to the raw, unnormalized output values produced before applying a final activation function like sigmoid or softmax. These logits represent the log-probabilities for each pixel or data point in the reconstructed output. The actual reconstruction likelihood *p*_*θ*_(**x**| **z**) would be computed from these logits using the specified distribution (Bernoulli for binarized or discretized logistic model for grayscale images, see A.3). In such a way, VAE is able to learn both mean (expected) value and the underlying uncertainty (variability).

##### A.3. Discretised logistic distribution

Images stored as 8-bit data contain discrete pixel values *k* ∈ {0, …, 255 }. To model these probabilistically with a neural decoder that outputs continuous parameters, a common choice is a discretised logistic distribution: a continuous logistic density is used only as an auxiliary object from which probability mass is assigned to each discrete pixel value. The decoder predicts a mean *µ* and a log-scale log *s* for each pixel, with *s* = exp(log *s*).

The discretised logistic cumulative distribution [44] function is:

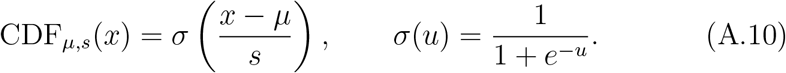

The probability mass assigned to pixel value *k* is the probability that a logistic random variable with parameters (*µ, s*) falls inside that bin:

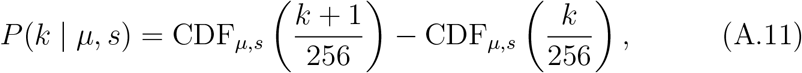

which is equivalently

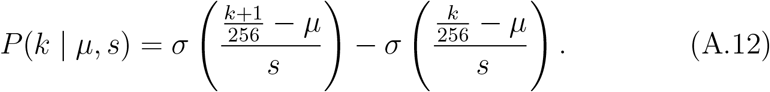

This construction converts a continuous distribution into a discrete probability mass function over the 256 possible pixel values. The log-likelihood of observing pixel value *k* is log *P* (*k µ, s*) and the reconstruction loss used in training is the negative log-likelihood, summed (or averaged) over all pixels.

#### A.4. Clamping encoder’s and decoder’s log-scale

As the model rapidly learns to reconstruct its input with great accuracy, reconstruction errors shrink and the estimated variance tends toward zero. [10] Rather than fixing the variance at a constant value, we clamp it using the HardTanh activation function, ensuring that the log-scale of each pixel remains within a reasonable range. [26]

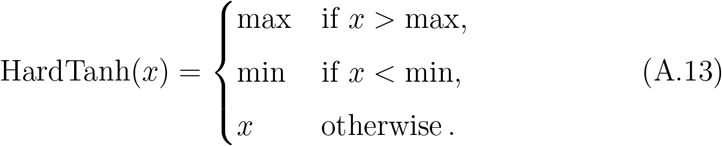

Using HardTanh activation function serves several purposes: *(1)* constraining the output prevents extreme values that could destabilize training or cause numerical issues; *(2)* limiting the output acts as a form of regularization, helping the model generalize better by avoiding overfitting to extreme values; and *(3)* in certain applications, especially in variational inference or generative models, controlling the output range can lead to more stable training dynamics.

#### B. Importance Weighted Autoencoder (IWAE)

Instead of a single draw from *q*_*ϕ*_(**z**| **x**), an ImportanceWeighted Autoen-coder (IWAE) [23, 24] is a VAE that tightens the ELBO by drawing *K* samples from the approximate posterior *q*_*ϕ*_(**z** |**x**). Each sample contributes importance weights that reflect how well it explains the input **x**.

In that way we replace the single Monte-Carlo estimate of log *p*_*θ*_(**x**) with a logarithm of the sum of exponentials (LogSumExp function, LSE) over *K* importance weights, which gives a tighter lower bound on log *p*_*θ*_(**x**) than the standard VAE ELBO as *K* increases.

Let’s start with drawing *K* independent latent samples:

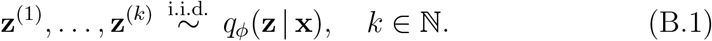

Since the encoder outputs only the mean *µ*_*ϕ*_(**x**_*i*_) and variance 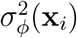 of the approximate posterior *q*_*ϕ*_(**z x**_*i*_), to generate actual latent samples 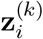 that simulate variability, we must sample using the reparameterization trick:

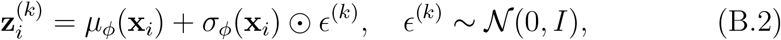

This introduces stochasticity needed for diverse reconstructions and training stability. Without sampling, there would be no variability — just a deterministic mapping. The unnormalised importance weight for the *k*-th sample is:

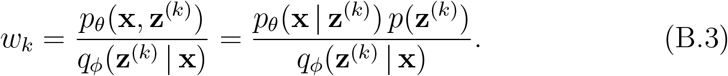

Each importance weight *w*_*k*_ quantifies how well sample **z**^(*k*)^ explains **x**, relative to the model’s proposal distribution *q*_*ϕ*_(**z** | **x**). The joint distribution *p*_*θ*_(**x, z**) is decomposed as *p*_*θ*_(**x** | **z**) *p*(**z**), where *p*_*θ*_(**x** | **z**) is the likelihood and *p*(**z**) is the prior. In other words, an importance weight *w*_*k*_ indicates how much the joint density *p*_*θ*_(**x, z**) deviates from the proposal density *q*_*ϕ*_(**z** | **x**). Consequently, the IWAE objective draws *K* samples from the approximate posterior *q*_*ϕ*_(**z** |**x**) to construct an importance-weighted estimate of the marginal likelihood, with the weights determined by the ratio of the joint distribution to the proposal distribution. [23]

When we require a probability distribution over the *K* samples (e.g., to compute gradients that involve a weighted average) we normalize importance weights as follows:

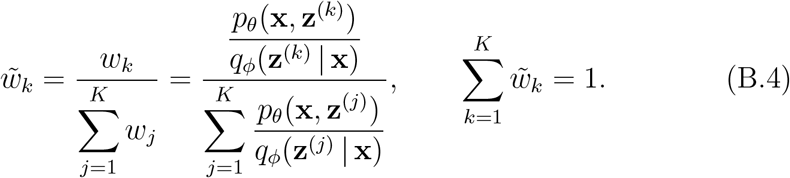

Therefore, IWAE objective for a single data point is the LSE (i.e., not just a simple average) of these weights:

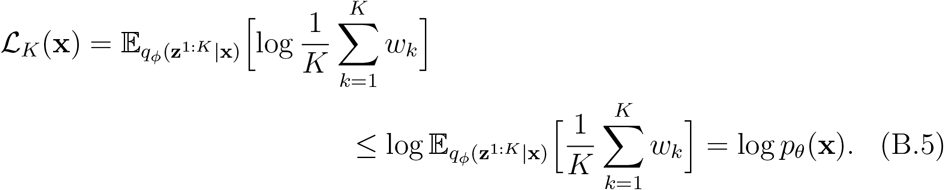

The inequality is strict in general — it’s the exact marginal likelihood, which the IWAE bound approaches as *K* → ∞. This allows the model to explore a richer set of latent configurations per datapoint. The bound also satisfies that ℝ_1_(**x**) reduces to the standard VAE ELBO.

In practice we approximate the expectation by drawing one set of *K* samples per data point. The so-called “log-sum-exp trick” refers to evaluating this expression in a numerically stable way. Each importance weight *w*_*k*_ = *p*_*θ*_(**x, z**^(*k*)^)*/q*_*ϕ*_(**z**^(*k*)^ | **x**) can be expressed as exp (log *p*_*θ*_(**x, z**^(*k*)^)−log *q*_*ϕ*_(**z**^(*k*)^ | **x**)). Hence the average weight 1*/K* ∑ *w*_*k*_ is a sum of exponentials. The IWAE objective contains the logarithm of this average, which is a LSE operation.

For a single latent sample **z**^(*k*)^ drawn from the variational posterior, the importance weight is:

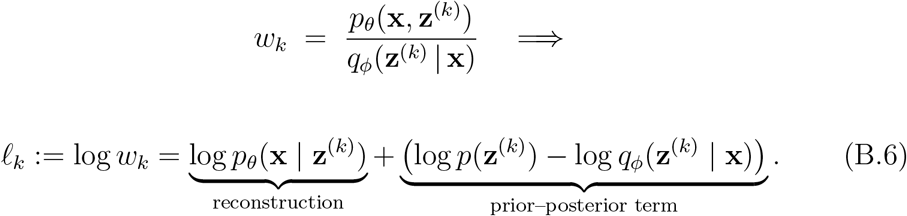

With *K* independent draws {**z**^(1)^, …, **z**^(*K*)^} the IWAE bound for a single data point is:

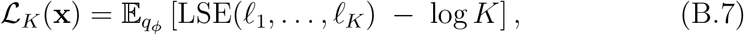

where the LSE function is computed in a numerically stable way as:

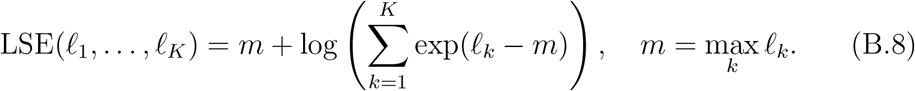

For a dataset 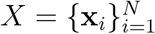 the overall objective is simply the average:

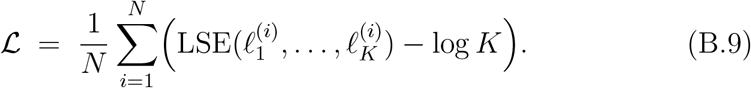

IWAE offers a more flexible approach by considering multiple samples from the approximate posterior *q*_*ϕ*_(**z** |**x**). The IWAE objective essentially encourages the model to find latent configurations that consistently explain the data well across these multiple samples. By averaging the importance weights (after applying the log-sum-exp trick), we effectively estimate the marginal likelihood using a weighted average of the likelihoods associated with each latent configuration. This weighted average can be seen as approximating the ELBO, but with a more nuanced approach to handling uncertainty in the latent space and providing stronger evidence for latent variables that explain the data well. [24]

Essentially, IWAE allows the model to explore a richer set of latent configurations per datapoint compared to standard VAEs, leading to better approximations of the marginal likelihood. Nevertheless, it is worth noting that the choice of *K* can significantly impact the accuracy and stability of the IWAE approximation.

### C. VAEs with Gaussian mixture distributions

#### C.1. Details and comments on VampPrior

To remind ourselves, in a standard VAE the prior is fixed and, more importantly, independent of the data. For generative purposes we sample **z** ∼ 𝒩 (0, *I*) and decode 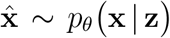. While simple, this approach often leads to sub-optimal generations if the aggregate posterior deviates significantly from the standard normal prior — a common issue known as *posterior collapse* or *poor latent coverage*. In particular, a simple prior (e.g. standard normal) can “over-regularise” the model by pulling all *q*_*ϕ*_ (**z** |**x**) toward one mode, leaving “holes” in the latent space where no data is represented.

With VampPrior the prior becomes a mixture of variational posteriors conditioned on learnable pseudo-inputs 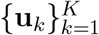 :

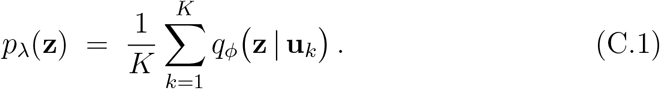

Each component *q*_*ϕ*_ (**z** | **u**_*k*_) is a Gaussian distribution output by the encoder for a pseudo-input **u**_*k*_. Since these pseudo-inputs are trainable parameters that are optimised during training, consequently the prior becomes multimodal and better captures the structure of the latent space, yielding more coherent and diverse samples. (see Supp. Fig. 1c.)

During training a real data point **x** is processed by the encoder to produce *q*_*ϕ*_ (**z** |**x**) ; this distribution is then sampled to obtain a latent code **z**, which the decoder uses to reconstruct the input as 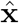. This reconstruction contributes to both the reconstruction loss and the KL divergence term in the ELBO. In contrast, the pseudo-inputs **u**_*k*_ are synthetic data points that are only passed through the encoder to shape the components of the prior; they are not decoded, so they contribute only to the KL term (no reconstruction loss).

According to Tomczak and Welling [26] the objective function for single datapoint **x**_∗_ using *L* Monte Carlo sample points is:

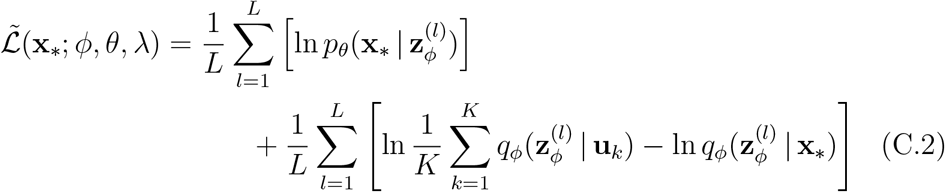

This design allows the model to learn a flexible, data-driven prior that better matches the aggregated posterior, thereby reducing over-regularisation. By choosing *K*≪ N pseudo-inputs we efficiently approximate the optimal prior without overfitting, which typically improves the ELBO and yields higher-quality generated samples.

Gradient updates follow this division: the encoder parameters *ϕ* receive gradients from both real data **x** and pseudo-inputs **u**_*k*_; the decoder parameters *θ* are updated only via real data (since only real inputs are recon-structed). The pseudo-inputs themselves are optimised through the KL term, allowing them to adaptively refine the prior structure over time. Hence, the training loop is essentially a standard VAE with an additional encoder pass over the *K* pseudo-inputs to construct the adaptive prior.

Critically, the pseudo-inputs **u**_*k*_ (which live in the same space as real inputs **x**) are learnable parameters. They are typically initialized randomly and optimised jointly with all network weights via back-propagation. Intuitively each **u**_*k*_ behaves like a prototype data point — the encoder produces a Gaussian in latent space for it, and these mixture components shape the overall prior to match the data. As Tomczak & Welling explain, the pseudo-inputs “mimic observable variables (e.g. images) and are learned by back-propagation”, making the prior rich and multimodal. [26]

#### C.2. Details and comments on Exemplar VAE

The Exemplar VAE replaces the standard VAE’s fixed isotropic Gaussian prior 𝒩 (0, *I*) with a data-driven, non-parametric prior (inspired by the VampPrior framework) formed as a mixture of latent encodings from actual training exemplars. [27] This enables more semantically meaningful and structured latent representations by leveraging real data points as references. (see Supp. Fig. 1d.)

In Exemplar VAE, there is a single encoder network, but it is applied to two types of inputs:

1. *q*_*ϕ*_(**z** |**x**) which is the variational posterior encoder used to encode an input **x** into the latent space, and
2. *r*_*ϕ*_(**z** | **x**_*n*_) which is the variational posterior encoder that maps each training exemplar **x**_*n*_ to a latent distribution, forming components of the prior mixture.

Hence, while the notation distinguishes *q*_*ϕ*_ and *r*_*ϕ*_, they represent the same parametric function evaluated at different inputs. The prior in the latent space is defined as a uniform mixture over encoded exemplars:

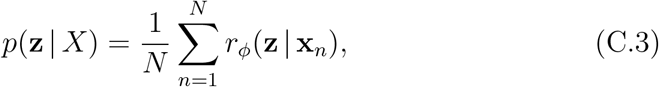

where the components *r*_*ϕ*_(**z** |**x**_*n*_) are learned Gaussians in the latent space, with both means and variances optimized during training.

Instead of minimizing *D*_KL_ (*q*_*ϕ*_(**z** |**x**) ∥ 𝒩 (0, *I*)), the Exemplar VAE minimizes:

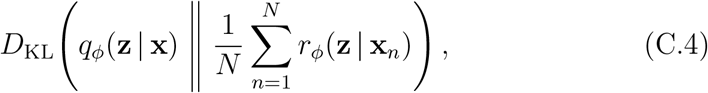

which encourages the variational posterior to align with the data-driven mixture prior. The log marginal likelihood for a data point **x** is:

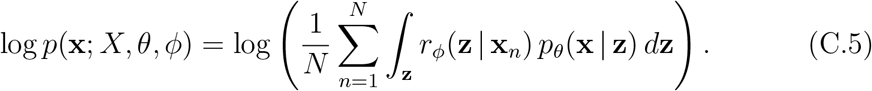

An evidence lower bound (ELBO) is derived as:

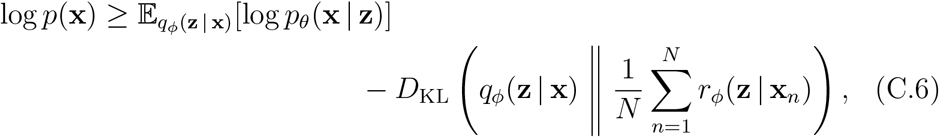

which is equivalently:

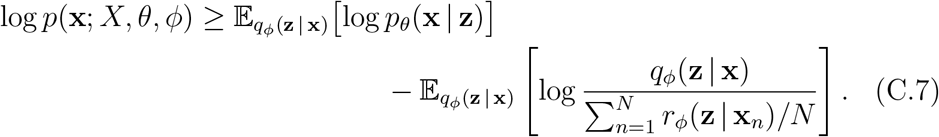

The encoder *r*_*ϕ*_(**z** |**x**_*n*_) is trained jointly with *q*_*ϕ*_(**z** |**x**) by optimizing the ELBO. Gradients from the KL divergence flow through the mixture components, enabling backpropagation into *r*_*ϕ*_. Thus, *r*_*ϕ*_ is updated implicitly via the ELBO, ensuring that the exemplar encodings adapt to better support the generative model.

To prevent overfitting and memorization authors used Exemplar Leave-One-Out and Exemplar Subsampling. However, the most important inovation to improve efficiency and tighten the ELBO was Retrieval-Augmented Training (RAT) that uses approximate nearest neighbor search in the latent space to select a subset of relevant exemplars for each input during training. This allows the model to focus on influential neighbors, improving generalization and generation quality.

And finally, new observations are generated by selecting an exemplar **x**_*n*_ and transforming it via a transition distribution defined by the encoder and decoder:

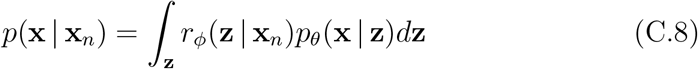

which reflects a generative mechanism rooted in real data transformations rather than purely synthetic sampling.

### D. Network architectures and training details

For this study, we utilized a collection of Variational Autoencoders (VAEs) implemented in PyTorch, building upon the VampPrior framework introduced by Tomczak and Welling [26], as detailed in Norouzi et al. [27]. The neural network layers consisted of fully-connected networks (MLPs), with layer dimensions specified within brackets using a dash to separate input and output units. [27] Below are the architectures of the encoder and decoder, respectively:

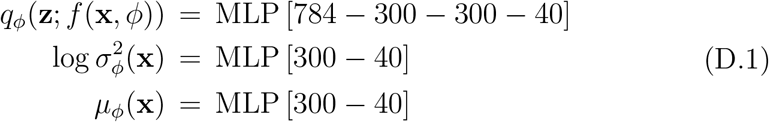

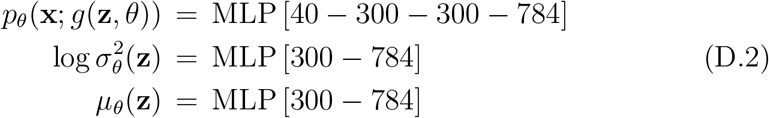

Tomczak and Welling [26] used the gating mechanism as an element-wise nonlinearity. Following their work, all models were trained for a maximum of 2,000 epochs with a mini-batch size of 100, a warm-up period of 100 epochs, early stopping with a look-ahead of 50 iterations, and 500 pseudo-inputs. We applied the Adam optimizer with normalized gradients for gradient descent with a learning rate of 5 ×10^−4^, *ϵ* = 1 ×10^−7^, *β*_1_ = 0.9, and *β*_2_ = 0.999; weights were initialized using the Glorot and Bengio method. (For more details, see their work [26], especially all iterations on arXiv.) These specifications establish a consistent baseline for evaluating the performance of VampPrior and Exemplar VAE, which have achieved state-of-the-art results on various datasets. [26]

### E. Definitions of evaluation metrics

For more information on evaluation metrics check the following references [1–3, 6].

The Adjusted Rand Index (ARI) measures the similarity between two clusterings by adjusting the Rand Index (RI) for chance agreement, providing a more reliable evaluation.

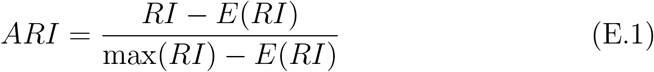

where:

- 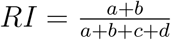: Raw Rand Index
  - *a*: Number of pairs in the same cluster in both true and predicted assignments.
  - *b*: Number of pairs in different clusters in both.
  - *c, d*: Number of pairs disagreeing on cluster membership.
- *E*(*RI*): Expected value of RI under random labeling. Alternatively, using a contingency table:

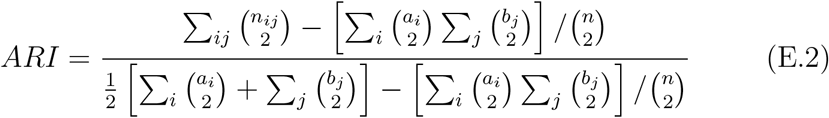

where:

- *n*_*ij*_: Number of samples in cluster *i* (true) and *j* (predicted).
- *a*_*i*_, *b*_*j*_: Marginal sums of the contingency table.

The Adjusted Mutual Information (AMI) measures the agreement between two clusterings by adjusting the Mutual Information (MI) score for chance, providing a more reliable comparison. Alternative normalization forms use arithmetic, geometric, or minimum entropy in the denominator.

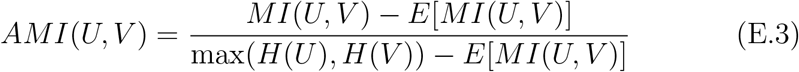

where:

- *MI*(*U, V*): Mutual Information between clusterings *U* and *V*.
- *E*[*MI*(*U, V*)]: Expected MI under random labeling with fixed cluster sizes.
- *H*(*U*), *H*(*V*): Entropies of the clusterings *U* and *V*.

The Fowlkes–Mallows Index (FMI) measures the similarity between two clusterings based on pairwise point relationships. It is calculated as:

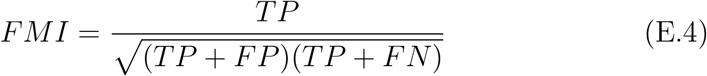

where TP, FP, and FN denote the number of true positives, false positives, and false negatives, respectively, when considering pairwise point cluster assignments.

The V–measure quantifies clustering quality by calculating the harmonic mean of homogeneity and completeness when comparing a predicted labeling to ground truth. It ranges from 0.0 to 1.0, with 1.0 indicating perfect homogeneity and completeness. The V-measure is equivalent to normalized mutual information using arithmetic averaging.

Homogeneity ensures each cluster contains only one class, while completeness guarantees all members of a class are in the same cluster. The formula is:

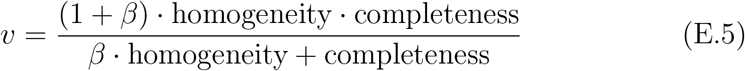

where *β* controls the weighting of completeness relative to homogeneity; *β >* 1 emphasizes completeness, and *β <* 1 emphasizes homogeneity. The default value is *β* = 1.0. The V–measure is symmetric and label-independent.

The Silhouette Score evaluates clustering quality by measuring how similar a data point is to its own cluster compared to others, based on cohesion and separation.

For each data point *i*:

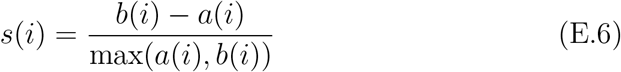

where:

- *a*(*i*) denotes the average distance to other points in the same cluster (cohesion).
- *b*(*i*) signifies the average distance to points in the nearest neighboring cluster (separation).

The overall Silhouette Score is the mean of *s*(*i*) across all data points.

The Davies–Bouldin Index (DBI) evaluates clustering quality by measuring the average similarity between each cluster and its most similar one, based on intra-cluster dispersion and inter-cluster separation.

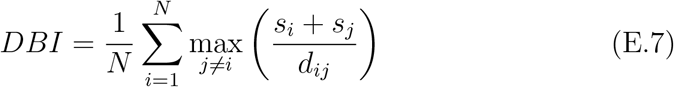

where:

- *N* represents the number of clusters.
- *s*_*i*_, *s*_*j*_ denote the average distance of points to their respective cluster centroids (intra-cluster dispersion).
- *d*_*ij*_ signifies the distance between the centroids of clusters *i* and *j* (intercluster separation).

The Calinski–Harabasz Index (CHI) evaluates clustering quality by measuring the ratio of between-cluster dispersion to within-cluster dispersion, adjusted for degrees of freedom.

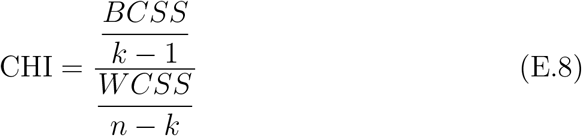

where:

- *BCSS*: Between-Cluster Sum of Squares (measures separation between clusters).
- *WCSS*: Within-Cluster Sum of Squares (measures compactness within clusters).
- *k*: Number of clusters.
- *n*: Total number of data points.

High silhouette scores coupled with low Davies–Bouldin values imply that the algorithm produces clusters that are compact, distinct, and ideally suited for applications beyond mere label matching. In other words, these metrics reveal whether the algorithm has identified useful cluster structures — even when labels are not perfectly recovered — making them essential for selecting the best clustering method rather than relying solely on accuracy.

### F. Dimensionality reduction and clustering

#### F.1. Comments on t-SNE and UMAP

The t-SNE algorithm first computes pairwise similarities between points in the high-dimensional space using a Gaussian kernel; it then models the low-dimensional space with a Student’s t-distribution. Finally, t-SNE minimizes the Kullback–Leibler divergence between these two probability distributions via gradient descent.

UMAP treats the dataset as a fuzzy topological manifold: first, it builds a weighted graph in the original space by connecting each point to its nearest neighbors; then it optimizes an embedding that keeps this fuzzy relation intact in the low-dimensional representation.

The authors of the UMAP algorithm set out to evaluate whether its embeddings are suitable for downstream clustering methods such as k-means and HDBSCAN, using the MNIST dataset as a benchmark. When comparing against plain PCA, they noted that UMAP yielded better cluster quality, attributing this advantage to its ability to capture strong manifold structure in the data. [29] In a similar fashion, An and Cho [9]’s VAE introduced *reconstruction probability* as a probabilistic anomaly score, which is a more principled and objective measure compared to the deterministic reconstruction error used by autoencoders and PCA-based methods.

#### F.2. Comments on k-means and HDBSCAN

K-means [30] is an unsupervised machine-learning algorithm that partitions a dataset into *K* distinct, non-overlapping clusters, each defined by a unique centroid. The algorithm iteratively updates the positions of these centroids to minimize the sum of squared distances between data points and their assigned cluster centers, ensuring that points within a cluster are as similar as possible while maintaining maximal separation between clusters.

Hierarchical Density-Based Spatial Clustering of Applications with Noise (HDBSCAN) [31] is an unsupervised clustering algorithm designed to identify clusters of varying densities and shapes within datasets while automatically determining the optimal number of clusters and identifying noise points. The HDBSCAN parameter that sets the minimum cluster size determines the smallest group considered a cluster; groups below this threshold are treated as noise and filtered out.

Accordingly, we evaluated clustering for three values: 50, 60, and 80. Our tests show that a minimum cluster size of 50 yields the best agreement with ground-truth labels on both t-SNE and UMAP embeddings while maintaining a comparable proportion of points assigned to clusters. We therefore adopt a fixed threshold of 60, which balances slightly increased coverage of clustered points against robust comparisons across VAE architectures and ensures that variations in ARI or accuracy reflect genuine latent-space grouping rather than the number of points discarded as noise.

#### F.3. Explanation of re-labelling procedures

The LOO-kNN supervised classification method can align predicted cluster labels with ground truth labels by employing a leave-one-out cross-validation strategy in combination with k-nearest neighbors (k-NN) classification. For each data point, that point is temporarily excluded from the dataset; the remaining points train a k-NN classifier that then predicts the label for the omitted point. Repeating this procedure for all data points constructs a mapping from true labels to cluster labels.

Heuristic labeling seeks to align predicted cluster labels with ground truth by assigning to each cluster the most frequent true class observed within it. Specifically, for every distinct cluster identified by the clustering algorithm, the method determines the predominant true class among its members and assigns that class label to all points in the cluster. The resulting output is a re-labelled clustering where cluster identifiers have been replaced with mapped true classes.

## References

[1] A. E. Ezugwu, A. M. Ikotun, O. O. Oyelade, L. Abualigah, J. O. Agushaka, C. I. Eke, A. A. Akinyelu, A comprehensive survey of clustering algorithms: State-of-the-art machine learning applications, taxonomy, challenges, and future research prospects, Engineering Applications of Artificial Intelligence 110 (2022) 104743. doi:10.1016/j.engappai.2022.104743.

[2] S. Pitafi, T. Anwar, Z. Sharif, A Taxonomy of Machine Learning Clustering Algorithms, Challenges, and Future Realms, Applied Sciences 13 (2023) 3529. doi:10.3390/app13063529.

[3] A. A. Wani, Comprehensive analysis of clustering algorithms: Exploring limitations and innovative solutions, PeerJ Computer Science 10 (2024) e2286. doi:10.7717/peerj-cs.2286.

[4] M. C. Thrun, Distance-based clustering challenges for unbiased bench-marking studies, Scientific Reports 11 (2021) 18988. doi:10.1038/s41598-021-98126-1.

[5] Q. Ji, Y. Sun, J. Gao, Y. Hu, B. Yin, A Decoder-Free Variational Deep Embedding for Unsupervised Clustering, IEEE Transactions on Neural Networks and Learning Systems 33 (2022) 5681–5693. doi:10.1109/TNNLS.2021.3071275.

[6] B. Roellinger, F. Thenier, C. Leclech, C. Coirault, E. Angelini, A. I. Barakat, Clustering cell nuclei on microgrooves for disease diagnosis using deep learning, Scientific Reports 15 (2025) 22476. doi:10.1038/s41598-025-05788-2.

[7] L. Huang, S. Ruan, Y. Xing, M. Feng, A review of uncertainty quantification in medical image analysis: Probabilistic and non-probabilistic methods, Medical Image Analysis 97 (2024) 103223. doi:10.1016/j.media.2024.103223.

[8] M. R. Karim, O. Beyan, A. Zappa, I. G. Costa, D. Rebholz-Schuhmann,M. Cochez, S. Decker, Deep learning-based clustering approaches for bioinformatics, Briefings in Bioinformatics 22 (2021) 393–415. doi:10.1093/bib/bbz170.

[9] J. An, S. Cho, Variational Autoencoder based Anomaly Detection using Reconstruction Probability, 2015.

[10] H. Akrami, A. A. Joshi, S. Aydore, R. M. Leahy, Addressing Variance Shrinkage in Variational Autoencoders using Quantile Regression, 2020. doi:10.48550/ARXIV.2010.09042.

[11] H. Akrami, A. Joshi, S. Aydore, R. Leahy, Quantile Regression for Uncertainty Estimation in VAEs with Applications to Brain Lesion Detection, in: A. Feragen, S. Sommer, J. Schnabel, M. Nielsen (Eds.), Information Processing in Medical Imaging, volume 12729, Springer International Publishing, Cham, 2021, pp. 689–700. doi:10.1007/978-3-030-78191-0_53.

[12] H. Akrami, A. A. Joshi, S. AydÖore, R. M. Leahy, Deep Quantile Regression for Uncertainty Estimation in Unsupervised and Supervised Lesion Detection, The Journal of Machine Learning for Biomedical Imaging 1 (2022) 008.

[13] J. Schmidhuber, Deep learning in neural networks: An overview, Neural Networks 61 (2015) 85–117. doi:10.1016/j.neunet.2014.09.003.

[14] I. Goodfellow, Y. Bengio, A. Courville, Deep Learning, MIT Press, 2016.

[15] D. P. Kingma, M. Welling, Auto-Encoding Variational Bayes, 2013. doi:10.48550/ARXIV.1312.6114.

[16] D. P. Kingma, M. Welling, An Introduction to Variational Autoen-coders, Foundations and Trends® in Machine Learning 12 (2019) 307–392. doi:10.1561/2200000056.

[17] J. D. Havtorn, J. Frellsen, S. Hauberg, L. Maaløe, Hierarchical VAEs Know What They Don’t Know, 2021. doi:10.48550/ARXIV.2102.08248.

[18] H. Xu, Y. Feng, J. Chen, Z. Wang, H. Qiao, W. Chen, N. Zhao, Z. Li, J. Bu, Z. Li, Y. Liu, Y. Zhao, D. Pei, Unsupervised Anomaly Detection via Variational Auto-Encoder for Seasonal KPIs in Web Applications, n: Proceedings of the 2018 World Wide Web Conference on World Wide Web-WWW ‘18, ACM Press, Lyon, France, 2018, pp. 187–196. doi:10.1145/3178876.3185996.

[19] Y. Guo, W. Liao, Q. Wang, L. Yu, T. Ji, P. Li, Multidimensional Time Series Anomaly Detection: A GRU-based Gaussian Mixture Variational Autoencoder Approach, in: J. Zhu, I. Takeuchi (Eds.), Proceedings of The 10th Asian Conference on Machine Learning, volume 95 of Proceedings of Machine Learning Research, PMLR, 2018, pp. 97–112.

[20] H. Khalid, S. S. Woo, OC-FakeDect: Classifying Deepfakes Using One-Class Variational Autoencoder, in: Proceedings of the IEEE/CVF Conference on Computer Vision and Pattern Recognition (CVPR) Work-shops, 2020.

[21] C.-C. M. Yeh, Y. Zhu, E. E. Papalexakis, A. Mueen, E. Keogh, Representation Learning by Reconstructing Neighborhoods, 2018. doi:10.48550/ARXIV.1811.01557.

[22] J. Guo, G. Liu, Y. Zuo, J. Wu, An Anomaly Detection Framework Based on Autoencoder and Nearest Neighbor, in: 2018 15th International Conference on Service Systems and Service Management (ICSSSM), IEEE, Hangzhou, China, 2018, pp. 1–6. doi:10.1109/ICSSSM.2018.8464983.

[23] Y. Burda, R. Grosse, R. Salakhutdinov, Importance Weighted Autoen-coders, 2015. doi:10.48550/ARXIV.1509.00519.

[24] C. Cremer, Q. Morris, D. Duvenaud, Reinterpreting Importance-Weighted Autoencoders, 2017. doi:10.48550/ARXIV.1704.02916.

[25] V. Prasad, D. Das, B. Bhowmick, Variational Clustering: Leveraging Variational Autoencoders for Image Clustering (2020). doi:10.48550/ARXIV.2005.04613.

[26] J. M. Tomczak, M. Welling, VAE with a VampPrior, 2017. doi:10.48550/ARXIV.1705.07120.

[27] S. Norouzi, D. J. Fleet, M. Norouzi, Exemplar VAE: Linking Generative Models, Nearest Neighbor Retrieval, and Data Augmentation, in:H. Larochelle, M. Ranzato, R. Hadsell, M. F. Balcan, H. Lin (Eds.), Advances in Neural Information Processing Systems, volume 33, Curran Associates, Inc., 2020, pp. 8753–8764.

[28] G. Hinton, S. Roweis, Stochastic neighbor embedding, in: Proceedings of the 16th International Conference on Neural Information Processing Systems, NIPS’02, MIT Press, Cambridge, MA, USA, 2002, pp. 857–864.

[29] L. McInnes, J. Healy, J. Melville, UMAP: Uniform Manifold Approximation and Projection for Dimension Reduction, 2018. doi:10.48550/ARXIV.1802.03426.

[30] S. Lloyd, Least squares quantization in PCM, IEEE Transactions on Information Theory 28 (1982) 129–137. doi:10.1109/TIT.1982.1056489.

[31] R. J. G. B. Campello, D. Moulavi, J. Sander, Density-Based Clustering Based on Hierarchical Density Estimates, in: D. Hutchison, T. Kanade,J. Kittler, J. M. Kleinberg, F. Mattern, J. C. Mitchell, M. Naor, O. Nier-strasz, C. Pandu Rangan, B. Steffen, M. Sudan, D. Terzopoulos, D. Tygar, M. Y. Vardi, G. Weikum, J. Pei, V. S. Tseng, L. Cao, H. Motoda, G. Xu (Eds.), Advances in Knowledge Discovery and Data Mining, volume 7819, Springer Berlin Heidelberg, Berlin, Heidelberg, 2013, pp. 160–172. doi:10.1007/978-3-642-37456-2_14.

[32] T. Iqbal, S. Qureshi, Reconstruction probability-based anomaly detection using variational auto-encoders, International Journal of Computers and Applications 45 (2023) 231–237. doi:10.1080/1206212X.2022.2143026.

[33] D. P. Gomari, A. Schweickart, L. Cerchietti, E. Paietta, H. Fernandez,H. Al-Amin, K. Suhre, J. Krumsiek, Variational autoencoders learn transferrable representations of metabolomics data, Communications Biology 5 (2022) 645. doi:10.1038/s42003-022-03579-3.

[34] C. Xu, Y. Dai, R. Lin, S. Wang, Deep clustering by maximizing mutual information in variational auto-encoder, Knowledge-Based Systems 205 (2020) 106260. doi:10.1016/j.knosys.2020.106260.

[35] M. Sabokrou, M. Khalooei, E. Adeli, Self-Supervised Representation Learning via Neighborhood-Relational Encoding, in: Proceedings of the IEEE/CVF International Conference on Computer Vision (ICCV), 2019.

[36] Y. Lee, H. Kwon, F. Park, Neighborhood Reconstructing Autoen-coders, in: M. Ranzato, A. Beygelzimer, Y. Dauphin, P. S. Liang, J. W. Vaughan (Eds.), Advances in Neural Information Processing Systems, volume 34, Curran Associates, Inc., 2021, pp. 536–546.

[37] D. Zimmerer, S. A. A. Kohl, J. Petersen, F. Isensee, K. H. Maier-Hein, Context-encoding Variational Autoencoder for Unsupervised Anomaly Detection, 2018. doi:10.48550/ARXIV.1812.05941.

[38] A. Vahdat, J. Kautz, NVAE: A Deep Hierarchical Variational Autoen-coder, in: H. Larochelle, M. Ranzato, R. Hadsell, M. F. Balcan, H. Lin (Eds.), Advances in Neural Information Processing Systems, volume 33, Curran Associates, Inc., 2020, pp. 19667–19679.

[39] C. K. Sønderby, T. Raiko, L. Maaløe, S. K. Sønderby, O. Winther, Ladder Variational Autoencoders, in: D. Lee, M. Sugiyama, U. Luxburg, I. Guyon, R. Garnett (Eds.), Advances in Neural Information Processing Systems, volume 29, Curran Associates, Inc., 2016.

[40] M. Tschannen, O. Bachem, M. Lucic, Recent Advances in Autoencoder-Based Representation Learning, 2018. doi:10.48550/ARXIV.1812.05069.

[41] M. J. Johnson, D. Duvenaud, A. B. Wiltschko, S. R. Datta, R. P. Adams, Structured VAEs: Composing graphical models with neural networks for structured representations and fast inference, 2017. doi:10.48550/arXiv.1603.06277. arXiv:1603.06277.

[42] E. Dupont, Learning Disentangled Joint Continuous and Discrete Representations, 2018. doi:10.48550/ARXIV.1804.00104.

[43] A. van den Oord, O. Vinyals, K. Kavukcuoglu, Neural Discrete Representation Learning, 2017. doi:10.48550/ARXIV.1711.00937.

[44] D. P. Kingma, T. Salimans, R. Jozefowicz, X. Chen, I. Sutskever,M. Welling, Improving Variational Inference with Inverse Autoregressive Flow, 2016. doi:10.48550/ARXIV.1606.04934.

